# Embodied Dynamic Causal Modelling: Brain and Cardiac Effective Connectivity

**DOI:** 10.64898/2026.10.27.762241

**Authors:** Andrew Levy, Peter Zeidman, Micah Allen, Karl Friston

## Abstract

Brain-body interactions underpin embodied inference and interoception, and are central to both general health and the pathophysiology of mental health disorders. These interactions are orchestrated across a hierarchy extending from the cortex, through the brainstem, to peripheral autonomic effectors. Existing methods either reduce this communication to scalar summaries or lack an invertible biophysical model, and so cannot resolve where in this extended architecture an observed effect originates. To address this, we introduce Embodied Dynamic Causal Modelling, an extension of the generative causal modelling framework that formalises hierarchical message passing across the full brain-body axis. We instantiate the framework for brain-heart interactions with the Brainstem Cardiac Network, a closed-loop neural mass model of brainstem autonomic circuitry and sinoatrial node dynamics. By incorporating five brainstem nuclei, a viscerosensory thalamic relay and the cardiac pacemaker, the network enables estimation of directed effective connectivity across the cortical-brainstem-cardiac axis.

As an illustrative case study, representative of the experiments and extended physiological systems that Embodied DCM and the BCN can now interrogate, we applied the framework to simultaneously recorded electroencephalography and electrocardiography during an auditory Go/No-Go paradigm. A multimodal observation model permits joint inversion of electrophysiological and cardiac responses, so that neural and bodily measurements together constrain an account of central and peripheral dynamics. The case study shows how the framework can arbitrate between competing hypotheses about functional cytoarchitecture, distinguishing granular from agranular microcircuits within the viscero-sensorimotor hierarchy. It also shows how time-varying estimates of synaptic efficacy can be recovered alongside continuous latent neuronal dynamics, over extended windows spanning anticipation, target processing and motor inhibition. By linking neural and cardiac signals to a single generative architecture, this work moves the study of brain-heart interactions beyond correlation, providing a framework for testing theories of embodied inference and for identifying circuit-level disruptions in disorders of interoceptive and autonomic regulation.

## Introduction

Cardiac-brain interactions are orchestrated across a hierarchy that extends from the intrinsic cardiac nervous system and spinal cord, through the brainstem and subcortical structures, to cortical regions (Ardell and Armour 2016). Communication along this axis is bidirectional, with descending autonomic commands setting physiological reference points at each level and ascending visceral afferents relaying sensory information back to higher processing centres (K. Friston 2010; Seth and Friston 2016). The traditional view casts the regulatory objective as the simple matching of cardiac output to metabolic demand (van Weperen and Vaseghi 2026), yet empirical findings routinely demonstrate that cardiac output frequently dissociates from metabolic need (Guyton, Coleman, and Granger 1972; Stöhr 2022) and instead reflects a multi-objective form of regulation encompassing sensory optimisation, threat avoidance, and social engagement (Barrett, Quigley, and Hamilton 2016; Skora, Livermore, and Roelofs 2022). This broad repertoire of goals finds a natural home within predictive processing and active inference, where descending commands are specified in terms of predictions of the expected physiological states for each subordinate tier of the hierarchy. The scope of these predictions broadens as one ascends from reflexive arcs in the periphery, which operate on fast timescales to manage immediate homeostatic perturbations, through subcortical structures such as the hypothalamus and amygdala, onwards to primary interoceptive regions such as the insula and anterior cingulate. And finally, to higher-order prefrontal regions, which generate predictions across the slowest behavioural timescales to incorporate allostatic goals, anticipatory adjustments that prepare the body for future needs rather than merely responding to current demands (Kiebel, Daunizeau, and Friston 2008; Barrett and Simmons 2015; Allen, Varga, and Heck 2023).

This hierarchical architecture has been the subject of numerous theoretical proposals concerning the various goal-oriented regimes it serves and its mapping onto the underlying functional and computational anatomy (Barrett, Quigley, and Hamilton 2016; Skora, Livermore, and Roelofs 2022; van Weperen and Vaseghi 2026; Allen et al. 2022). Nevertheless, these frameworks remain largely untested empirically. This is largely due to shortcomings in the predominant methods used to characterise brain-heart interactions. Existing approaches typically either reduce the system to a scalar summary or capture only a partial aspect of its bidirectional communication. For example, heart rate variability (HRV), the most widely deployed index, collapses the integrated output of non-linear interactions across multiple temporal scales into a univariate observable (Bill-man 2011). The result is a profound many-to-one degeneracy, whereby a given HRV value can arise from qualitatively distinct physiological configurations, including altered sympathovagal balance, changes in baroreflex gain, respiratory pattern, or intrinsic cardiac dynamics, with no means of disambiguation from the index alone (Billman 2013; Hayano and Yuda 2021). Alternatively, directed statistical measures such as Granger causality and transfer entropy partially address this by attempting to recover causal asymmetries between neural and cardiac time series. However, without an explicit generative model of the underlying physiology, the dependencies they recover cannot be distinguished from spurious coupling arising from shared inputs or unmeasured confounding sources (Karl J. Friston et al. 2014; Faes, Porta, and Nollo 2015).

Recent methodologies include a forward model of peripheral cardiac dynamics, with some integrating this bidirectionally with neural activity to assess brain-heart interactions (Catrambone et al. 2019; Candia-Rivera 2023). However, the implicit architecture is not readily amenable to inversion, thus precluding the recovery of latent autonomic states from the observed signals. A related approach involves principled Bayesian inversion of a generative model of cardiac dynamics, which yields a full posterior distribution over latent heart rate trajectories (Rosas et al. 2024). However, this generative model is phenomenological rather than biophysical, and lacks any representation of brain-heart interactions. The core limitation shared by all these approaches is the absence of an invertible generative biophysical model that maps the full hierarchical, circular causal architecture onto measured signals. Consequently, the level of the hierarchy at which an observed coupling change originates, whether peripheral, brainstem, or cortical, cannot be identified from the data.

Dynamic Causal Modelling (DCM) offers a principled solution to these limitations. The DCM framework rests on two key innovations. First, DCM inverts a biophysically plausible generative model against observed data, employing a formal separation between a hidden neuronal state space and a modality-specific observation model that maps those hidden states onto observed signals (K. J. Friston, Harrison, and Penny 2003; David et al. 2006). Second, the use of variational Bayesian inversion provides an approximation to the model evidence, which allows competing hypotheses about the causal architecture of the hidden state space to be evaluated through Bayesian model comparison (BMC) (Penny et al. 2004). DCM addresses the many-to-one degeneracy, inherent in conventional brain-heart indices, by making the mapping from hidden neural states to observables explicit through the forward model. DCM has been applied extensively to characterise directed effective connectivity within cortical networks, but has not previously been extended to incorporate the brainstem autonomic circuitry and peripheral cardiac effectors. As a result, effective connectivity estimates have been confined to cortical sources, and the descending and ascending pathways through which the brain commands and monitors the heart have remained outside the scope of generative inference.

Here, we introduce the Brainstem Cardiac Network (BCN), a closed-loop neural mass model of cardioautonomic brainstem circuitry and sinoatrial node (SAN) dynamics, designed for integration into the DCM framework to model event-related responses. The BCN embeds five key brainstem nuclei, the periaqueductal gray (PAG), parabrachial nucleus (PBN), nucleus tractus solitarius (NTS), nucleus ambiguus (NA), and rostral ventrolateral medulla (RVLM), a viscerosensory thalamic nucleus, and the SAN as the terminal peripheral cardiac effector. Deployment of the BCN alongside existing cortical and subcortical microcircuits enables estimation of directed effective connectivity across the full cortical-brainstem-cardiac axis. Achieved through a single joint inversion of multimodal electrophysiological and cardiac data.

In the present paper, we detail the architecture of the BCN and motivate each component within an active inference account of cardiac regulation. We then demonstrate the model with an empirical case study applying DCM to simultaneously recorded EEG and ECG during an auditory Go/No-Go task, a paradigm known to elicit dissociable cardiac responses across anticipation, target processing, and motor inhibition (Ribeiro and Castelo-Branco 2019). To our knowledge, this is the first unified generative model to estimate directed effective connectivity across the visceromotor cortical-brainstem-cardiac axis and to jointly invert multimodal neural and cardiac data. The illustrative case study focused on (i) validation in relation to known functional anatomy and (ii) showcasing the ability to characterise fluctuations in neuronal activity and synaptic efficacy.

## Methods

In this section, we introduce a biophysical model of brainstem cardiac control designed for inversion within the DCM framework. The BCN enables characterisation of bidirectional communication among cortical and subcortical regions and the brainstem, supporting hypothesis testing of the functional architectures underlying heart rate (HR) modulation. The circuitry is intentionally constrained to provide a generalisable model of short-timescale HR control relevant to cognitive and affective paradigms. Physical exertion and metabolic demand also modulate HR through many of the same autonomic pathways represented in the BCN. However, they introduce additional regulatory pathways that are not included in the present model. The BCN is therefore intended to explain transient HR modulation under conditions of relatively stable metabolic demand, rather than the broader cardiovascular adjustments accompanying sustained exertion or changing metabolic requirements.

We first detail the neurophysiological motivation for the inclusion of each node and its connectivity, and assign a tentative active (interoceptive) inference-inspired computational role in cardiac regulation. We then describe the multimodal observation model that enables joint inversion from simultaneously acquired EEG and ECG data. Finally, we present an empirical case study applying the BCN within the DCM for Sequential Responses (DCM-SR) framework to an auditory Go/No-Go task (A. D. Levy, Zeidman, and Friston 2026a).

### Brainstem Cardiac Network

The BCN comprises five brainstem nuclei, a viscerosensory thalamic nucleus, and the SAN (Figure 1). These nodes collectively instantiate the functional architecture through which descending visceromotor predictions and ascending viscerosensory signals are mediated. The PAG arbitrates descending sympathovagal policy; the NTS provides viscerosensory state estimation informing parasympathetic outflow; the PBN and thalamic nucleus form the ascending interoceptive relay; and the NA and RVLM serve as the parasympathetic and sympathetic actuators, respectively, with the SAN as the peripheral effector where chronotropic drives are resolved. Extrinsic descending visceromotor commands (i.e. setpoints) from cortical and subcortical sources terminate on the PAG and NTS, whilst ascending viscerosensory signals are relayed via the PBN and thalamus to cortical regions, completing the interoceptive feedback loop.

**Figure 1.**
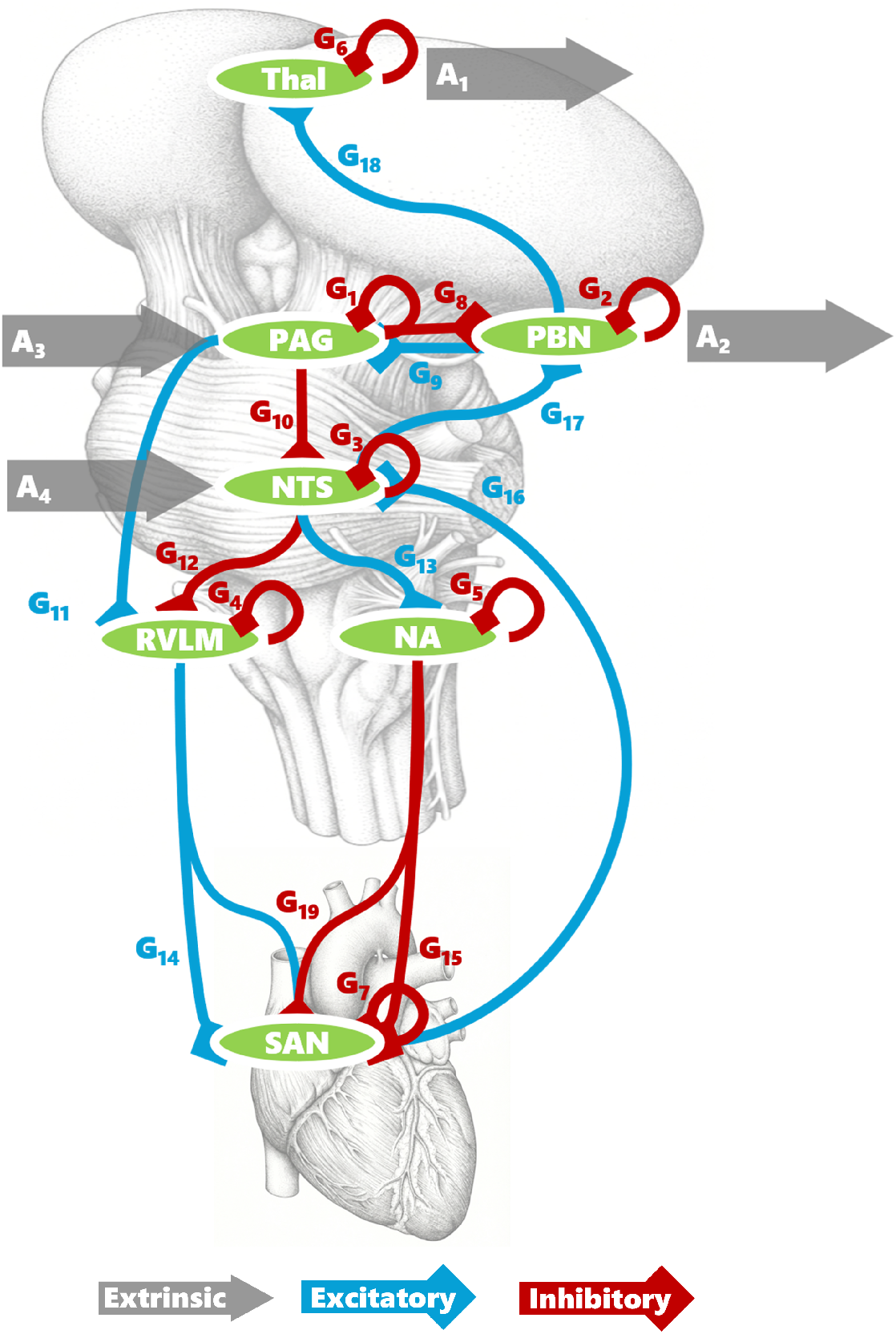
Schematic of the Brainstem Cardiac Network (BCN): The network comprises five primary brainstem regions: Periaqueductal Gray (PAG), Parabrachial Nucleus (PBN), Nucleus Tractus Solitarius (NTS), Rostral Ventrolateral Medulla (RVLM), and Nucleus Ambiguus (NA), which interact to regulate the Sinoatrial Node (SAN) and a viscerosensory Thalamic nucleus (Thal). Excitatory and inhibitory intrinsic connections are shown in blue and red, respectively; red circular arrows denote intrinsic self-inhibition. Extrinsic connections: *A*_1_ denotes ascending output from the thalamus; *A*_2_ denotes ascending output from the PBN to the cortex; *A*_3_ denotes descending input to the PAG; and *A*_4_ denotes descending input to the NTS.

The BCN is designed primarily to test predictive processing and active inference accounts of cardiac regulation, in which brainstem nuclei implement phylogenetically conserved homeostatic reflexes that are driven by ascending viscerosensory prediction errors and overridden by descending visceromotor commands from higher-order structures (Pezzulo, Parr, and Friston 2021; Allen 2020). The neurocomputational roles assigned to each node below are presented as testable hypotheses to guide future inquiry rather than as a definitive computational architecture.

### Brainstem Nuclei

Each brainstem nucleus in the BCN is modelled as a neural mass whose equations of motion take the same generic form as cell populations in the DCM canonical microcircuit (CMC; Bastos et al. (2012)). The use of the same mathematical form as the CMC ensures that the BCN is fully interoperable with existing DCM architectures and that parameters carry consistent biophysical interpretations across cortical and subcortical regions.

Each population is equipped with two states: a membrane potential, *x_v_*, and a synaptic current, *x_c_*. Presynaptic firing is expressed as a deviation from baseline via a sigmoid activation function

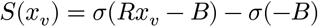

where *R* is the gain of the activation function and *B* is the bias term setting the resting operating point. The total synaptic input to a population is then

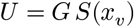

where *G* is the signed intrinsic connection strengths, with positive values denoting excitatory and negative values denoting inhibitory influences. Convolution of this pre-synaptic input with the synaptic response function produces post-synaptic depolarisation, described by the second-order ordinary differential equation

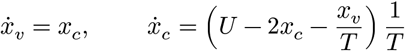

where *T* is the synaptic time constant governing the temporal evolution of the postsynaptic response.

In what follows, we detail the specific parameterisation of each nucleus in terms of its synaptic time constant *T*, intrinsic gain *R*, and afferent connection strengths *G*, which together fully characterise the equations of motion for that population (Table 1). These values are best regarded as physiologically motivated priors rather than definitive estimates. The current data are insufficient to refine the sympathetic-parasympathetic kinetic separation because the short epochs and single paradigm used limit the ability to disambiguate the two based on their time constants. Confidently establishing these constants requires multiple datasets, paradigms, and targeted perturbations that differentially engage the limbs. We treat the kinetic asymmetry as an open question, noting that the framework is well suited to address it as more data become available.

**Table 1.** Brainstem Cardiac Network Prior Parameters.

| Category | Parameter | Value | Description / Region |
|---|---|---|---|
| **Time Constants (*T*)** | *T*<sub>1</sub> | 4 ms | Periaqueductal Gray (PAG) |
|  | *T*<sub>2</sub> | 4 ms | Parabrachial Nucleus (PBN) |
|  | *T*<sub>3</sub> | 4 ms | Nucleus Tractus Solitarius (NTS) |
|  | *T*<sub>4</sub> | 192 ms | Rostral Ventrolateral Medulla (RVLM) |
|  | *T*<sub>5</sub> | 32 ms | Nucleus Ambiguus (NA) |
|  | *T*<sub>6</sub> | 384 ms | Sinoatrial Node (SAN) |
|  | *T*<sub>7</sub> | 4 ms | Thalamus (THAL) |
|  | *T*<sub>8</sub> | 0.1 ms | Heart Rate (HR) |
| **Sigmoid Gains (*R*)** | *R*<sub>*n*</sub> | 2/3 | Neuronal activation gain |
|  | *R*<sub>*a*</sub> | 3/2 | Arousal activation gain |
|  | *R*<sub>*h*</sub> | 1/4 | Heart-rate activation gain |
| **Sigmoid Offset (*B*)** | *B*<sub>*n*</sub> | 0 | Neuronal activation offset |
|  | *B*<sub>*a*</sub> | 2 | Arousal activation offset |
|  | *B*<sub>*h*</sub> | 0 | Heart-rate activation offset |
| **Intrinsic Connections (*G*)** | *G*<sub>1</sub> | 800 | PAG Self-Inhibition |
|  | *G*<sub>2</sub> | 800 | PBN Self-Inhibition |
|  | *G*<sub>3</sub> | 400 | NTS Self-Inhibition |
|  | *G*<sub>4</sub> | 400 | RVLM Self-Inhibition |
|  | *G*<sub>5</sub> | 400 | NA Self-Inhibition |
|  | *G*<sub>6</sub> | 800 | THAL Self-Inhibition |
|  | *G*<sub>7</sub> | 25 | SAN Self-Inhibition |
|  | *G*<sub>8</sub> | 400 | PAG → PBN |
|  | *G*<sub>9</sub> | 200 | PBN → PAG |
|  | *G*<sub>10</sub> | 400 | PAG → NTS |
|  | *G*<sub>11</sub> | 200 | PAG → RVLM |
|  | *G*<sub>12</sub> | 400 | NTS → RVLM |
|  | *G*<sub>13</sub> | 1200 | NTS → NA |
|  | *G*<sub>14</sub> | 400 | RVLM → SAN |
|  | *G*<sub>15</sub> | 800 | NA → SAN |
|  | *G*<sub>16</sub> | 2 | SAN → NTS |
|  | *G*<sub>17</sub> | 400 | NTS → PBN |
|  | *G*<sub>18</sub> | 800 | PBN → THAL |
|  | *G*<sub>19</sub> | 1200 | RVLM × NA → SAN (nonlinear) |

#### Periaqueductal Gray (Phylogenetic Cardiac Policy Selection)

The PAG serves as a key integrative hub of autonomic functions and survival-related behaviours, acting as an interface between forebrain emotional systems and lower brainstem effectors (Behbehani 1995; Linnman et al. 2012). It is organised into four longitudinal columns, each coordinating a distinct autonomic-behavioural programme (Bandler and Shipley 1994). The dorsolateral column mediates anticipatory anxiety and threat assessment (R. A. L. Dampney et al. 2013), the dorsomedial column regulates innate aversion and respiratory drive with a central role in panic and breathlessness (Faull and Pattinson 2017), the lateral column drives active coping characterised by flight, hypertension, and tachycardia, and the ventrolateral column orchestrates passive coping manifesting as freezing, hypotension and bradycardia (Rossi, Maione, and Berrino 1994). The PAG columns are under tonic GABAergic inhibition and mutually inhibitory, such that intrinsic lateral inhibition resolves columnar competition into a single dominant output (Tovote et al. 2016). This positions the PAG as a plausible site for cardiac policy selection (Benarroch 2012).

In the BCN, the intrinsic connection priors favour vagal output, setting a default parasympathetic policy of cardiac deceleration. Recruitment of the PAG provides the principal pathway for overriding this default and engaging a sympathetic tachycardic policy, arising through increased descending extrinsic input (*A*_3_), increased inhibitory projection from the PAG to NTS (*G*_10_) releasing the vagal brake, and increased excitatory projection from the PAG to the RVLM (*G*_11_) driving sympathetic outflow. Sympathetic withdrawal and deepening cardiac deceleration follow from reductions in these same projections. The BCN also captures passive coping via the reciprocal PAG to PBN projection, discussed below.

#### Nucleus Tractus Solitarius (State Estimation under Parasympathetic Policies)

The NTS is a critical integrative node where descending commands from forebrain and diencephalic structures are assimilated with ascending visceral interoceptive afference (Forstenpointner et al. 2022). High-fidelity afference from arterial baroreceptors and chemoreceptors arrives via the glossopharyngeal and vagus nerves, providing a composite representation of cardiovascular and respiratory states. Whilst descending projections from the insula and infralimbic cortices, the central amygdala, and the bed nucleus of the stria terminalis synapse directly onto the NTS, providing a substrate through which cortically elaborated allostatic expectations can bias autonomic reflex gain (R. A. Dampney 1994; Casar, Idiaquez, and Benarroch 2025).

The NTS can be viewed as a state estimator serving parasympathetic policies with goals of homeostatic conservation. Deviations from physiological setpoints are treated as (precision-weighted) prediction errors, which the NTS attempts to minimise via local vagal adjustments. When prediction errors are small or persistent, resolution is attempted within the prevailing vagal policy. However, when they are large or sustained, inadequacy of the current parasympathetic policy is propagated as an ascending alarm signal to the PBN.

In the BCN, the NTS serves as the principal node by which the default vagal policy is selected or withdrawn. Selection is achieved by increased descending extrinsic input (*A*_4_), representing the forebrain’s allostatic endorsement of the default vagal regime; increased excitatory NTS to NA connectivity (*G*_13_) to augment vagal outflow; and increased inhibitory NTS to RVLM connectivity (*G*_12_), collapsing the polysynaptic relay via the caudal ventrolateral medulla into a single effective parameter suppressing sympathetic drive. The NTS receives ascending cardiac afference from the SAN (*G*_16_) as a crude approximation of baroreceptor signalling, constituting the principal means by which cardiac state is represented within the model.

#### Parabrachial Nucleus (State Estimation under Sympathetic Policies)

The PBN serves a dual role, both relaying ascending sensory information and arbitrating homeostatic threat. As a principal relay hub, it integrates ascending afferents from NTS, spinal dorsal horn, and trigeminal dorsal horn neurons; including interoceptive signals related to taste, pain, and autonomic control as well as exteroceptive inputs (R. A. Dampney 1994; Pauli et al. 2022; Chiang et al. 2019). Under non-threatening conditions, this multimodal information is conveyed principally via the parabrachial-thalamic projection, providing the thalamus with the integrated visceral and somatosensory context required for (possibly conscious) interoceptive inference (A. D. Craig 2002).

A parallel threat-detection role is mediated by a distinct PBN subpopulation sensitive to homeostatic disturbances that exceed the vagally mediated resolution capacity (Palmiter 2018; Pyeon et al. 2025). Upon detecting irreducible prediction errors, the PBN first attempts local resolution through influencing PAG policy selection (Bandler et al. 2000; Hayward 2007). When threat magnitude precludes local resolution, direct projections bypass the thalamic relay to reach subcortical and cortical structures (Grady et al. 2020), positioning the PBN as the state estimator for defensive sympathetic policies and enabling alarm escalation beyond brainstem arbitration.

In the BCN, ascending NTS to PBN signals (*G*_17_) convey interoceptive information that, at sufficient connection strength, signals disturbances beyond vagal resolution. These drive an excitatory PBN to PAG projection (*G*_9_), enforcing sympathetic dominance until cortical policy updates. Conversely, PAG states inhibit the PBN (*G*_8_), suppressing alarm signalling and consolidating the bradypnoea and autonomic quiescence of the freeze state. The PBN transmits ascending signals to the thalamus (*G*_18_) and directly to the cortex via extrinsic output (*A*_2_), the latter accounting for direct PBN afferents to deep pyramidal cells of granular and dysgranular interoceptive regions (Grady et al. 2020). We note that in the current implementation these connections remain operative when linking the BCN to agranular microcircuit regions, despite the relative sparsity of direct PBN innervation in those areas.

#### Parabrachial Nucleus (State Estimation under Sympathetic Policies)

The posterior ventromedial (VMpo) and ventral posterior parvocellular (VPpc) thalamic nuclei serve as the principal relay for ascending discriminative interoception, via dense afferents relayed through the PBN (A. Craig 2003; Jerath and Malani 2026). Under parasympathetic policies, this relay provides high-precision visceral feedback to the cortex. However, during defensive mobilisation, PAG inhibition of the NTS withdraws the discriminative interoceptive stream, preventing homeostatic restoration signals from competing with the active coping policy (Lovick 1992; Nosaka et al. 1993).

Within the BCN, the thalamus receives ascending sensory data from the PBN (*G*_18_), formalising the NTS-PBN-thalamus relay as the canonical ascending interoceptive sensory channel (Forstenpointner et al. 2022; A. D. Craig 2002). The ascending output (*A*_1_) represents thalamocortical transmission to the spiny stellate population of the posterior insula, consistent with the laminar and regional specificity of VMpo and VPpc projections.

When connecting the BCN to agranular regions such as the anterior insula, a direct thalamic projection targets the middle pyramidal cells. This is a simplification, as insular afference is mediated principally via intra-insular forward connections from the posterior insula; a dedicated model of the insular cytoarchitectural gradient is a natural direction for future work (Allen 2020).

#### Nucleus Ambiguus (Parasympathetic Action)

The NA contains preganglionic motor neurons that project via myelinated vagal fibres to postganglionic neurons within the intrinsic cardiac ganglia, innervating the SAN to provide a high-fidelity, beat-to-beat constraint of pacemaker rate (Gourine et al. 2016; McAllen and Spyer 1978). Under baseline conditions, tonic excitatory drive from the NTS sustains metabolic conservation (Neff, Mihalevich, and Mendelowitz 1998; Huang, Wang, and Ran 2025); during defensive mobilisation, PAG-mediated suppression of the NTS withdraws this drive, releasing the vagus brake and permitting cardiac acceleration (Farkas, Jansen, and Loewy 1997).

Within the BCN, the NA is modelled as the final brainstem output stage of the parasympathetic hierarchy, receiving strong excitatory drive from the NTS (*G*_13_) to instantiate the vagus brake. The descending connection from the NA to the SAN (*G*_15_) represents the peripheral vagal pathway, in which myelinated preganglionic B-fibres synapse on postganglionic neurons within the intrinsic cardiac ganglia and acetylcholine release onto muscarinic receptors at the SAN translates the central parasympathetic policy into a negative chronotropic effect.

#### Rostral Ventrolateral Medulla (Sympathetic Action)

The RVLM and the rostral raphe pallidus (rRPa) constitute the two principal sympathetic premotor nuclei of the medulla, governing functionally distinct limbs of the cardiovascular defence response. The RVLM is the final common pathway for sympathetic vasomotor tone, tonically inhibited by the NTS-mediated baroreflex but overridden during threat processing by descending PAG drive (Guyenet 2006; R. A. L. Dampney 2016). The rRPa mediates stress-evoked cardiac acceleration, tonically suppressed by GABAergic inhibition but disinhibited during psychological and physiological challenge to drive tachycardia via cardiac sympathetic preganglionic neurons in the intermediolateral cell column (Cao and Morrison 2003; S. F. Morrison and Gebber 1982). Despite their anatomical segregation, both nuclei are recruited concurrently by shared descending drive from the PAG and dorsomedial hypothalamus, producing coordinated vasoconstriction and tachycardia as the integrated cardiovascular defence response (Moraes et al. 2020; Horiuchi et al. 2004).

To reduce model complexity, we collapse the RVLM and rRPa into a single composite sympathetic premotor node, organised around RVLM-mediated baroreflex processes. We consider the baroreflex pathway the most relevant to cognitive and interoceptive paradigms. This lumping is justified by the functional co-linearity of the two nuclei under psychological load. However, the simplification breaks down in paradigms where they are dissociably recruited. For instance, cardiac-mediated thermoregulation performed by rRPa is preferentially engaged to drive tachycardia in the absence of RVLM-mediated vasoconstriction (Zaretsky, Zaretskaia, and DiMicco 2003; Shaun F. Morrison and Nakamura 2011).

Within the BCN, the RVLM node receives descending sympathoexcitatory drive from the PAG (*G*_11_), capturing the bulbospinal pathway through which central defence commands are translated into peripheral sympathetic outflow. The efferent connection to the SAN (*G*_14_) simplifies the polysynaptic pathway through the intermediolateral cell column and paravertebral sympathetic ganglia into a single effective parameter representing the net chronotropic drive produced by noradrenaline release onto beta-adrenergic receptors at the cardiac accelerator terminals.

### Cardiac Model

The cardiac model completes the BCN by resolving competing autonomic drives into chronotropic modulation of SAN firing rate. To do so, the SAN extends the standard neural mass formulation with a first-order HR state variable for observable inversion and an asymmetric bivariate activation function with a bilinear interaction term to capture nonlinear chronotropic effects. These extensions provide a minimal yet physiologically grounded description of chronotropic regulation compatible with DCM inversion.

#### Sinoatrial Node

The SAN is the principal cardiac pacemaker and the site where competing autonomic inputs are integrated to modulate HR. The relationship between autonomic inputs and chronotropic modulation departs from simple linear control in two respects. First, the system operates asymmetrically around a resting state held substantially below the intrinsic pacemaker rate by a dominant vagal tone. The two branches therefore have fundamentally different dynamic ranges: parasympathetic withdrawal can only raise HR as far as the intrinsic ceiling, whereas sympathetic excitation can drive it well beyond. Second, the two branches interact nonlinearly when coactivated, a phenomenon termed *accentuated antagonism*, whereby vagal efficacy is amplified in the presence of elevated sympathetic tone (M. N. Levy 1971; G. G. Berntson, Cacioppo, and Quigley 1991).

To capture these properties, we modify the SAN pre-synaptic activation function in two ways. First, the standard sigmoid *S* is replaced by an asymmetric variant *S_a_* for the RVLM and NA projections to the SAN:

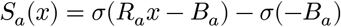

where *R_a_* = 3/2 and *B_a_* = 2. The positive bias *B_a_* positions the resting state in the lower tail of the sigmoid, such that negative perturbations rapidly saturate against a floor near −0.12 whilst positive perturbations can reach approximately +0.88, yielding a roughly sevenfold asymmetry in dynamic range.

Second, accentuated antagonism is captured by adding a bilinear interaction term to the net autonomic drive:

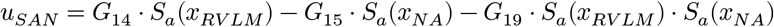

where *G*_14_ and *G*_15_ are the sympathetic and parasympathetic connection strengths, respectively. The third term, *G*_19_, is a bilinear product of both branches carrying a negative sign, such that concurrent activation yields a net inhibitory effect exceeding their linear sum, ensuring that vagal perturbations produce a larger bradycardic effect when sympathetic tone is elevated.

The resulting bivariate surface reproduces the four regimes of autonomic space (Fig. 2): high sympathetic with low parasympathetic drive yields strong positive net SAN drive, whilst the opposite produces strong negative net drive. Co-activation generates a net output shaped by accentuated antagonism, biasing the surface toward net negative chronotropy. Co-inhibition produces a net positive shift toward the intrinsic pacemaker rate, because tonic parasympathetic restraint dominates resting heart rate and its withdrawal outweighs the concurrent loss of sympathetic support (Gary G. Berntson, Cacioppo, and Quigley 1993; M. N. Levy 1971).

**Figure 2.**
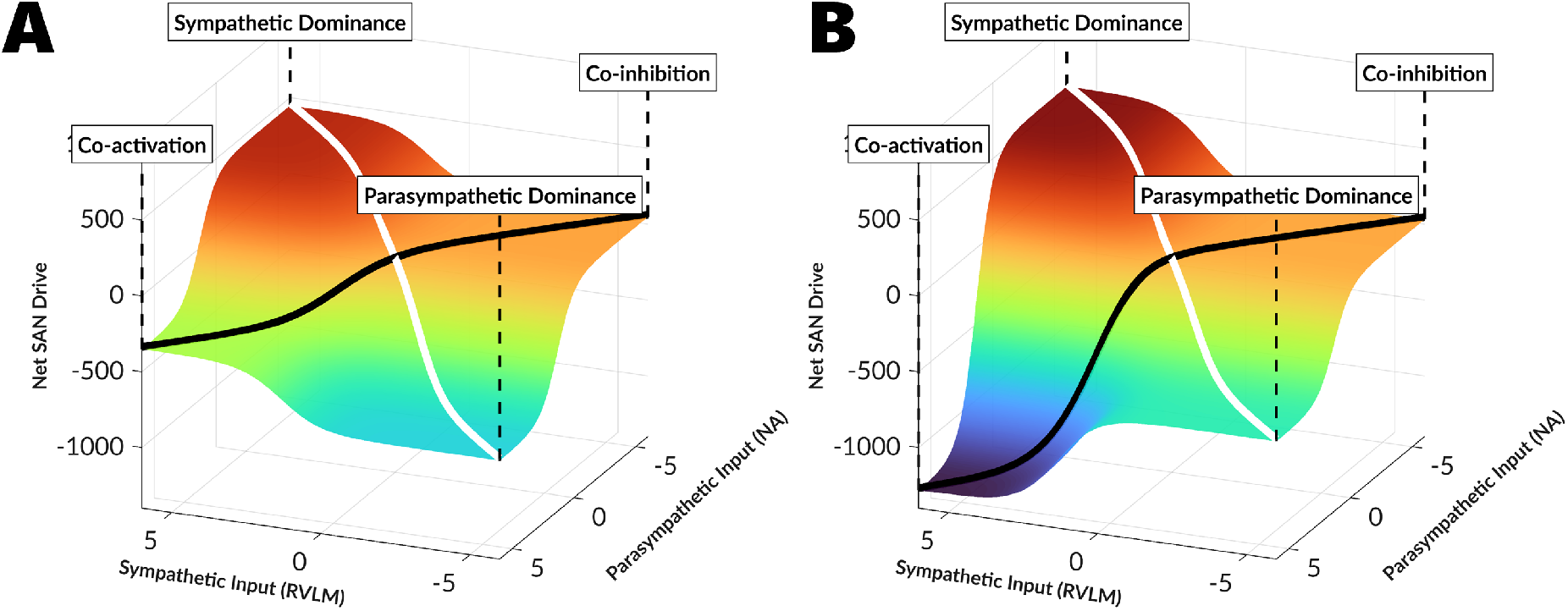
The Bivariate Autonomic Response Surface: illustration of net chronotropic drive to the sinoatrial node as a function of RVLM and NA inputs. Two model variants are shown: (A) linear SAN, in which the bilinear interaction term is omitted; and (B) nonlinear SAN, in which the bilinear term is included, capturing accentuated antagonism. By mapping these synaptic drives through an asymmetric sigmoid activation function, the model instantiates the bivariate autonomic space formalised by Berntson, Cacioppo, and Quigley (1991). The reciprocity axis (white line) captures the classical reciprocal model of autonomic control, extending between the poles of sympathetic and parasympathetic dominance. The coactivity axis (black line) captures the coactivity dimension, ranging from high-level coactivation to low-level coinhibition.

#### Heart Rate State Variable

In addition to the second-order membrane dynamics shared with the brainstem nuclei, the SAN includes a first-order heart rate state variable *x*_ℎ*τ*_:

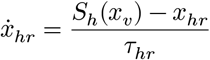

where *S*_ℎ_ is a shallow sigmoid (gain *R*_ℎ_ = 1/4) and no bias term, *x_v_* is SAN membrane potential, and *τ*_ℎ*τ*_ a short time constant (0.1 ms). This acts as a near-instantaneous transfer function, bounding SAN output for observation rather than representing a physiological process. In principle, lengthening *τ*_ℎ*τ*_ could crudely absorb measurement lags, such as those introduced by photoplethysmography rather than ECG. The transfer converts integrated SAN membrane voltage into a bounded HR variable, with the asymptotes of *S*_ℎ_ imposing upper and lower limits on *x*_ℎ*τ*_ that reflect the finite dynamic range of cardiac chronotropy across autonomic states. By representing this nonlinearity within the latent state dynamics rather than the observation function, *x*_ℎ*τ*_ can be related to the cardiac observation through a linear mapping, as required by the Bayesian inversion scheme described below.

### Observation Models

Standard DCM for ERPs maps a shared latent state space to a single neuroimaging modality. We extend the framework to permit multimodal observation mappings, in which a single hidden state vector is projected through modality-specific observation functions. Each function weights the contributions of distinct neural populations and hidden states independently and accounts for the unique gain and noise of its modality. Inversion uses the spm_nlsi_N Bayesian (Variational Laplace) scheme, which constrains each observation function to be a linear mapping from hidden states to observables (Zeidman, Friston, and Parr 2023). This multimodal extension permits modalities that were previously inverted separately, such as EEG and MEG, to be inverted conjointly under a shared generative model. It also opens the framework to additional physiological measures, including respiration and pupillometry, provided a generative model of each modality can be specified. In this study, the mapping is bivariate, comprising an electrophysiological lead-field model and a cardiac observation model (COM), each of which is described below (see Table 2).

**Table 2.** Multimodal Observation Model Extension Functions.

| **Category** | **Function** | **Description** |
|---|---|---|
| **Observation Models** | spm_lx_seq | Observation model function caller |
|  | spm_lx_erp | Electrophysiological observation model |
|  | spm_lx_com | Cardiac observation model |
| **Prior Moments** | spm_L_erp_priors | Prior moments for the lead-field parameters of ERP models |
|  | spm_L_com_priors | Prior moments for observation parameters of COM models |

### Electrophysiological Observation Model

The electrophysiological observation model follows the standard DCM for evoked responses using the equivalent current dipole (ECD) formulation, mapping the hidden state vector *x* to observed scalp potentials *y_eτp_* via a conventional lead-field model *L*(*θ_L_*), whose parameters encode the position and orientation of ECD, and a parameter vector *J_eτp_* determining the relative contribution of individual neural states:

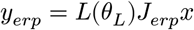

### Cardiac Observation Model

The COM linearly scales the HR state variable to the observed HR

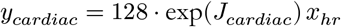

Here, *J_caτdiac_* is a log-scaling parameter that ensures the scaling factor remains positive, with the constant (128) setting the prior expected gain between the bounded model state and observed HR; the effective gain is then estimated from the data. This ensures preservation of sign such that increases in SAN membrane potential elevate HR and decreases reduce it.

### Observation Noise Model

In DCM for ERPs, observation noise is modelled using temporal covariance functions that capture residual autocorrelation, typically as an AR(1) process. For the multimodal data, we assigned independent AR(1) noise components to each modality, allowing modality-specific serial correlations.

### Empirical Case Study

The BCN is designed for deployment alongside existing cortical and subcortical models within the DCM framework. Doing so enables mechanistic inference over hierarchical cortical-brainstem-cardiac circuits, and permits joint modelling of visceromotor, interoceptive, exteroceptive, cognitive, and motor processes under a single generative model. This allows formal quantification of effective connectivity between central and cardiac systems. To illustrate these capabilities, we applied DCM-SR to an auditory Go/No-Go paradigm with simultaneous EEG-ECG, in which motor execution and inhibition unfold alongside measurable cardiac dynamics.

The analysis targeted three questions, framed primarily to validate the framework. First, we asked whether our framework could recover the functional role of the insula and mid-cingulate cortex within the interoceptive hierarchy, distinguishing a viscerosensory contribution, expressed through a granular microcircuit, from a visceromotor one, expressed through an agranular microcircuit. This constitutes an empirical test of theoretical proposals concerning the cytoarchitectural organisation of visceral sensorimotor cortex (Barrett and Simmons 2015). Second, we asked whether the data supported a nonlinear account of SAN innervation, specifically whether accentuated antagonism is necessary to explain the observed cardiac dynamics. Third, we sought to illustrate the capacity of the framework to resolve cardioceptive processing in terms of hierarchical message passing. We address this by interrogating the distinct contributions of rINS and lMCC to descending cardiac control and ascending interoceptive signalling, as well as establishing the identifiability of specific intrinsic brainstem connections mediating these effects.

### Empirical Dataset

We used an open-access auditory go/no-go dataset (https://openneuro.org/datasets/ds003690; Ribeiro and Castelo-Branco (2019)) comprising simultaneously acquired EEG and ECG from 36 participants (16 younger adults: 24 ± 3 years, 12 female; 20 older adults: 61 ± 6 years, 17 female) with normal or corrected vision and hearing and no neurological or psychiatric history. Each trial presented an anticipatory cue followed by either a go signal (250 ms, 1700 Hz tone, 80 trials/run) or a no-go signal (250 ms, 1300 Hz tone, 20 trials/run) at approximately 67 dB, requiring key presses for go trials and response withholding for no-go trials. Cue-to-target intervals ranged between 1.5 and 4.1s, whilst post-stimulus intervals ranged from 5.2 to 15.5s. EEG and ECG were recorded simultaneously at 500 Hz using a 64-channel Neuroscan system, with scalp electrodes placed according to the International 10-20 standard (reference CPz, ground FPz) and bipolar ECG electrodes placed on the chest. Each participant completed two approximately 8-minute runs. We analysed correctly performed go and no-go trials only, discarding behavioural errors and artefact-contaminated epochs.

### EEG Preprocessing

Continuous EEG data were imported into SPM25 (Wellcome Centre for Human Neuroimaging, London) and band-pass filtered using a fourth-order Butterworth filter (0.1-30 Hz), downsampled to 200 Hz, and epoched from −1000 to +6000 ms relative to trial onset. Epochs were rejected if any channel exceeded an absolute z-score of 7. Channels were re-referenced to the common average. Jittered cue-to-target intervals were corrected using Event-Related Warping (A. Levy, Zeidman, and Friston 2026), with onset cue and target events represented as Gaussians (SD = 50 ms) at the recorded stimulus times. Motor response jitter was not corrected to preserve temporal equivalence between go and no-go trials. Warping functions used eighth-order DST basis sets with Tukey tapering (taper ratio = 0.1) estimated over 18 interpolation steps. Trial-averaged ERPs were computed using unweighted averaging and grand-averaged across participants for DCM analysis.

### ECG Preprocessing

Continuous ECG data were band-pass filtered (8-40 Hz, fourth-order Butterworth) to isolate the QRS complex, with signal polarity corrected automatically to ensure positive R-peaks. QRS enhancement used the squared gradient of the signal smoothed with an 80 ms moving average, robustly Z-transformed via the median absolute deviation. R-peaks were detected using a local maxima algorithm (500 ms refractory period, prominence threshold 0.2). RR intervals were mapped to a continuous HR series in beats per minute via previous-neighbour interpolation. The same warping functions derived from ERW were applied to the HR data, preserving temporal alignment across modalities (A. Levy, Zeidman, and Friston 2026).

### DCM Specification

DCM-SR was applied to grand-averaged ERPs spanning −500 to +4000 ms relative to trial onset, partitioned into five sequential phases: onset cue, anticipation, motor preparation, target cue, and motor execution (Table 3). Baseline extrinsic connectivity (A), modulatory connectivity (B), and synaptic gains (G) were all treated as time-varying parameters. The BCN was embedded within a nine-source hierarchical cortical-brainstem network (Table 4), drawing on three neural mass models: CMC, motor microcircuit (MMC), and BCN. Bilateral auditory cortices (left and right superior temporal gyrus; lSTG, rSTG) received auditory inputs. A prefrontal control hierarchy (right dorsolateral prefrontal cortex, right inferior frontal gyrus, and left pre-supplementary motor area; rDLPFC, rIFG, lpSMA) arbitrated Go/No-Go task demands, with the rDLPFC receiving a slow cortical input during the anticipation phase and an ERP input during motor execution. The somatomotor system consisted solely of left primary motor cortex (lM1). The visceromotor system comprised two streams converging on the BCN: a cingulate-mediated stream (lpSMA to left mid-cingulate cortex; lMCC) and an insula-mediated stream (rIFG to right insula; rINS).

**Table 3.** DCM-SR Event Specification.

| Event | Onset<br>(ms) | Duration<br>(ms) | Bump<br>Function | Input<br>Node | Network<br>Spec |
|---|---|---|---|---|---|
| Onset Cue | 90 | 16 | spm_erp_u | rSTG & lSTG | Fig 3 A |
| Anticipation | 400 | 64 | spm_scp_u | rDLPFC | Fig 3 B |
| Motor Preparation | 1500 | 32 | spm_null_u | Nil | Fig 3 C |
| Target Cue | 1800 | 16 | spm_erp_u | rSTG & lSTG | Fig 3 D |
| Motor Execution | 2100 | 24 | spm_erp_u | rDLPFC | Fig 3 E |

**Table 4.** DCM-SR Source Specification.

| Source | NMM | MNI |  |  | Observation Model |
|---|---|---|---|---|---|
|  |  | x | y | z |  |
| rDLPFC | CMC | 30 | 43 | 23 | ECD |
| lMCC | CMC/MMC<sup>*</sup> | −6 | −10 | 40 | ECD |
| rINS | CMC/MMC<sup>*</sup> | 36 | 23 | −2 | ECD |
| rSTG | CMC | 60 | −22 | 8 | ECD |
| lSTG | CMC | −60 | −22 | 8 | ECD |
| lM1 | MMC | −38 | −14 | 46 | ECD |
| lpSMA | CMC | −6 | 6 | 58 | ECD |
| rIFG | CMC | 52 | 14 | 8 | ECD |
| Brainstem | BCN | Silent |  |  | COM |

<sup>\*</sup>rINS and lMCC were each assigned either a CMC or MMC architecture across separate inversions as part of a factorial model comparison; see Model Space.

We permitted experimental modulation of somatomotor reciprocal connections (lpSMA-rIFG, lpSMA-lM1, and rIFG-lM1) to explain motor inhibition during No-Go relative to Go trials. These modulations are not analysed further here and have been reported previously for this dataset (A. D. Levy, Zeidman, and Friston 2026a). For the visceromotor hierarchy, the system of primary interest, we modulated the extrinsic reciprocal connections between lMCC and the BCN and between rINS and the BCN. We permitted modulation of cortical intrinsic self-inhibition connections, specifically superficial pyramidal (SP) cells in the CMC and deep pyramidal (DP) cells in the MMC.

Within the brainstem, four connections were modulated: (1) PAG self-inhibition, included as a gain control over the selected sympathovagal policy; (2) NTS self-inhibition, included as a gain control over parasympathetic state estimation; (3) the descending PAG to NTS inhibitory connection accounting for modulation of the vagal brake; and (4) the PAG to RVLM connection, included as the direct sympathoexcitatory pathway.

### Model Space

We constructed a model space defined by two factors: (1) autonomic modulation of the sinoatrial node, specifically whether a bilinear term representing the modulatory effect of RVLM and NA activity on SAN firing rate was included (nonlinear SAN) or omitted (linear SAN); and (2) the cytoarchitecture of the two viscero-sensorimotor cortical sources, rINS and lMCC, which were assigned either CMC (granular) or MMC (agranular). All combinations were permitted, yielding eight models in total. Model comparison was performed using fixed-effects BMC based on the approximate log-model evidence. The winning model was defined as the one with the highest posterior probability, provided that this probability was at least 0.75.

### Bayesian Model Reduction

The winning model underwent Bayesian model reduction (BMR) to identify the most parsimonious functional anatomy underlying cardiac responses during the cue-target and motor-execution phases. BMR was applied directly to the posterior of the grand-averaged DCM, evaluating whether reduced models, in which task-modulatory parameters are fixed at zero, provide a better account of the data (Karl J. Friston et al. 2016; Zeidman et al. 2019).

Two separate BMRs were performed, one for each task phase of interest, each operating over the ten mod ulatory parameters summarised in Table 5, yielding a model space of 2_10_ = 1024 reduced models per phase. The first targeted the mechanisms underlying the transient cardiac acceleration observed during nogo cuetarget processing, asking whether this was mediated by descending visceromotor commands via the lMCC or rINS pathways, modulation of the vagal brake, sympathetic recruitment, or changes in gain at the level of the PAG or NTS. The second targeted the mechanisms underlying the deepening cardiac deceleration on no-go relative to go trials during motor execution, asking the same questions with respect to the slowing cardiac response. Where a reduced model attained a posterior probability exceeding 0.75, it was selected as the winning model; otherwise, parameter inference was conducted on the Bayesian model average (BMA), weighting estimates by their posterior probability across the reduced model space (Trujillo-Barreto, Aubert-Vázquez, and Valdés-Sosa 2004).

**Table 5.** Parameters subjected to Bayesian model reduction.

| Connection | Connection Type |
|---|---|
| rINS SP (self) | rINS Intrinsic |
| rINS → Bstem | Extrinsic |
| Bstem → rINS | Extrinsic |
| lMCC DP (self) | lMCC Intrinsic |
| lMCC → Bstem | Extrinsic |
| Bstem → lMCC | Extrinsic |
| PAG (self) | Bstem Intrinsic |
| NTS (self) | Bstem Intrinsic |
| PAG → NTS | Bstem Intrinsic |
| PAG → RVLM | Bstem Intrinsic |

### Sensitivity Analysis

The winning model was also subjected to a sensitivity analysis to characterise the ensuing changes in electrophysiological and cardiac responses consequent on perturbations to the parameters of interest. To aid interpretation, parameters were perturbed in the direction of their posterior effect. For the brainstem responses, sensitivity was assessed across all brainstem nuclei and the thalamus, summarised as the signed area under the curve accumulated over the full epoch window. For the HR responses, the full sensitivity timeseries was examined, preserving the temporal profile of the chronotropic response to each parameter perturbation. In addition, to characterise parameter identifiability across modalities, we computed the (numerical) derivative of the predicted outputs with respect to each parameter and subjected it to singular value decomposition (SVD), thereby quantifying each parameter’s contribution to observable variation in each modality.

## Results

In this section, we present the results of inverting the BCN within the DCM framework. We first tested whether a bilinear interaction between sympathetic and parasympathetic inputs at the SAN was necessary to account for HR modulation, and whether lMCC and rINS were better captured by granular or agranular circuitry motifs. The winning model was then advanced to characterise the functional architecture mediating cardiac responses to No-Go targets via BMR, over the viscero-sensorimotor and brainstem modulatory synaptic efficacies, followed by a sensitivity analysis that characterised how individual parameters contributed to latent brainstem responses and HR dynamics.

### Sinoatrial Node Nonlinearity and Viscero-sensorimotor Cytoarchitecture

We first addressed two questions: whether chronotropic modulation of the SAN required a non-linear term to capture accentuated antagonism (the potentiation of vagal slowing that arises when both autonomic branches are active); and whether the nominated viscero-sensorimotor regions (rINS and lMCC) were better represented as viscerosensory or visceromotor within the interoceptive hierarchy. The latter was operationalised as a comparison between two neural mass microcircuits: the CMC, whose granular structure reflects a predominantly viscerosensory role, and the MMC, whose agranular structure reflects a visceromotor role. To address these questions, we performed BMC across eight inverted DCMs, comprising all combinations of linear and non-linear SAN coupling and CMC or MMC representations of rINS and lMCC. The comparison strongly favoured Model 2, a linear SAN with an MMC (agranular) representation of lMCC and a CMC (granular) representation of rINS (*P_p_* > 0.99, Δ*F* = 21.39) (Figure 4A).

**Figure 3.**
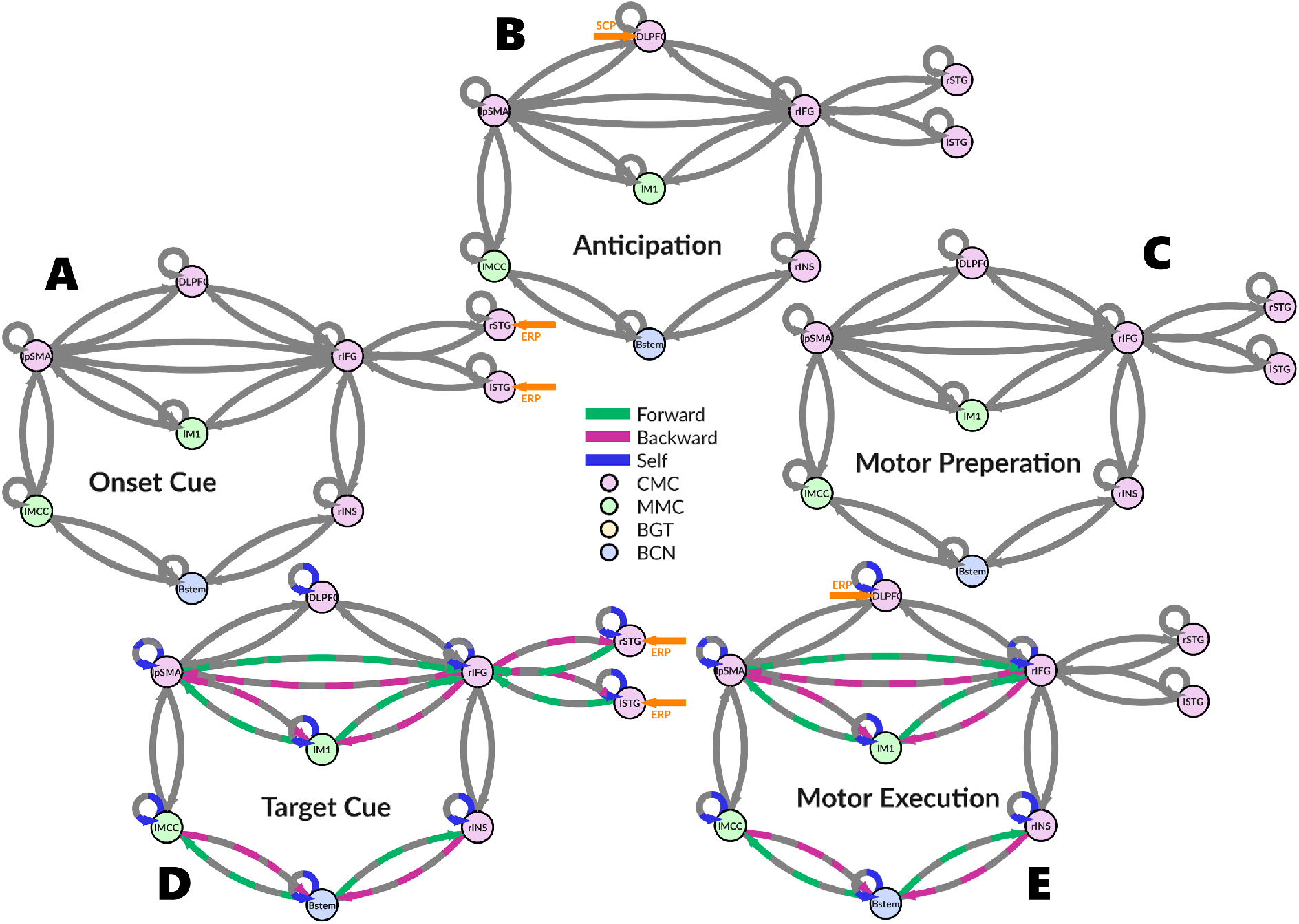
Network architecture and connectivity modulation across sequential events: Five panels (A–E) illustrate network configurations at onset cue (90 ms), anticipation (400 ms), motor preparation (1500 ms), target cue (1800 ms), and motor execution (2100 ms). Grey connections represent baseline extrinsic connectivity maintained throughout all events. Coloured connections indicate experimentally modulated connections. Orange arrows denote exogenous driving inputs modelled as bump functions for event-related potentials (ERP) and slow cortical potentials (SCP).

**Figure 4.**
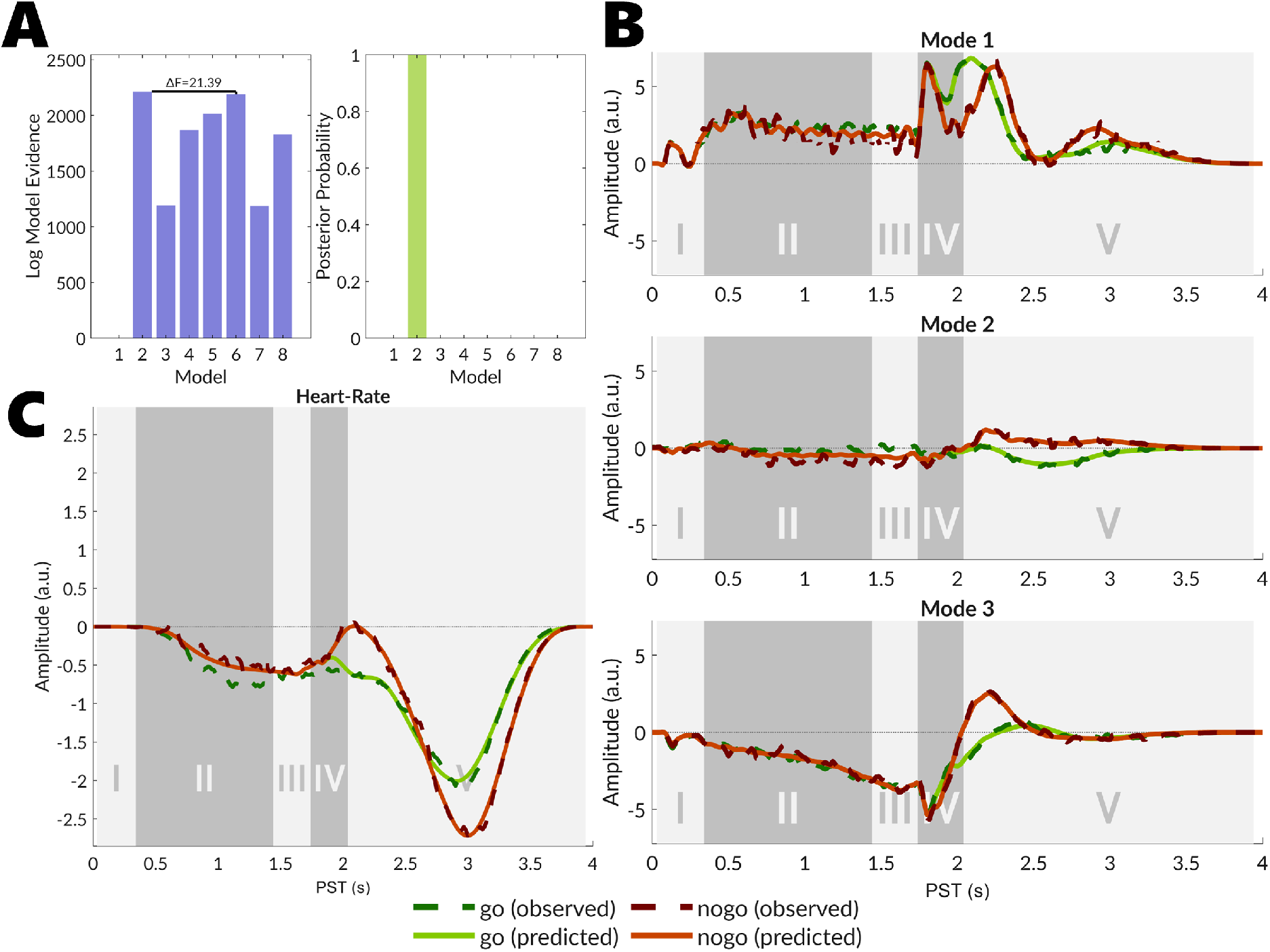
Bayesian model comparison and model fit. (A) Comparison across eight models varying SAN coupling (linear vs. nonlinear) and laminar cytoarchitecture for rINS and lMCC (CMC or MMC). Left: approximate log model evidence (free energy) relative to the lowest-performing model. Right: posterior probability, with model 2 winning decisively (>0.99). Model definitions: 1: Linear-SAN, rINS-CMC, lMCC-CMC; 2: Linear-SAN, rINS-CMC, lMCC-MMC; 3: Linear-SAN, rINS-MMC, lMCC-CMC; 4: Linear-SAN, rINS-MMC, lMCC-MMC; 5: Nonlinear-SAN, rINS-CMC, lMCC-CMC; 6: Nonlinear-SAN, rINS-CMC, lMCC-MMC; 7: Nonlinear-SAN, rINS-MMC, lMCC-CMC; 8: Nonlinear-SAN, rINS-MMC, lMCC-MMC. (B) Fit to the first three principal EEG modes. (C) Fit to heart rate timeseries. Grey shaded regions in (B) and (C) denote experimental phases: (I) onset cue (90-400 ms), (II) anticipation (400-1500 ms), (III) motor preparation (1500-1800 ms), (IV) target cue (1800-2100 ms), (V) motor execution (2100-4000 ms).

Figure 4 (B&C) displays the winning model fit to the first three EEG modes and HR timeseries for the winning DCM. Quantitative assessment across all nine EEG modes and HR yielded *R*_2_ = 96.46% for the go condition and *R*_2_ = 94.53% for the no-go condition, indicating that the DCM captures the large majority of systematic variance in both neural and cardiac signals.

### Onset Cue, Anticipation, Motor Preparation

Each trial began with an auditory onset cue (phase I), followed by an anticipatory window (phase II), and a motor preparation window (phase III), all immediately preceding the target presentation. Across these three phases, the observed HR showed a modest and sustained deceleration (Figure 4C). The recovered synaptic efficacy trajectories remained close to baseline, with no substantial fluctuations (Fig. 5), whilst the estimated membrane potentials demonstrated marked fluctuations (Fig. 6). The posterior estimates reported here are expressed as log-scaled deviations from their priors and are interpreted qualitatively. Additionally, given the BCN’s intrinsic bradycardic bias, only modest changes in connectivity were required to produce deceleration in response to descending command. As the synaptic efficacy parameters were largely unremarkable, we focus instead on the estimated neuronal states.

**Figure 5.**
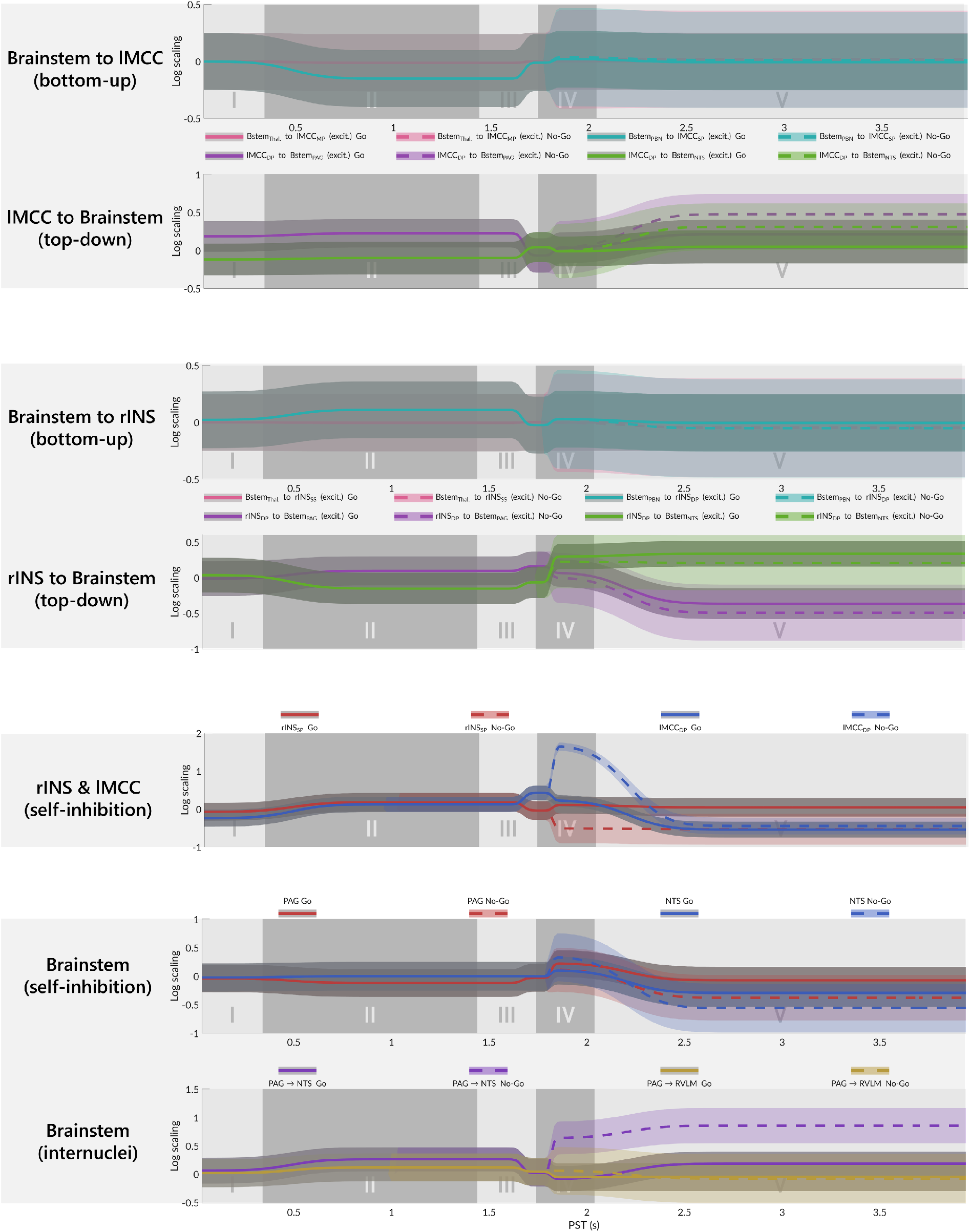
Parameter Trajectories: Time-varying posterior expectations (lines) and posterior standard deviations (shaded bands) for modulatory effects on extrinsic and intrinsic connections involving the brainstem. Grey shaded regions delineate the five sequential experimental phases: (I) onset cue (90-400 ms), (II) anticipation (400-1500 ms), (III) motor preparation (1500-1800 ms), (IV) target cue (1800-2100 ms), (V) motor execution (2100-4000 ms).

**Figure 6.**
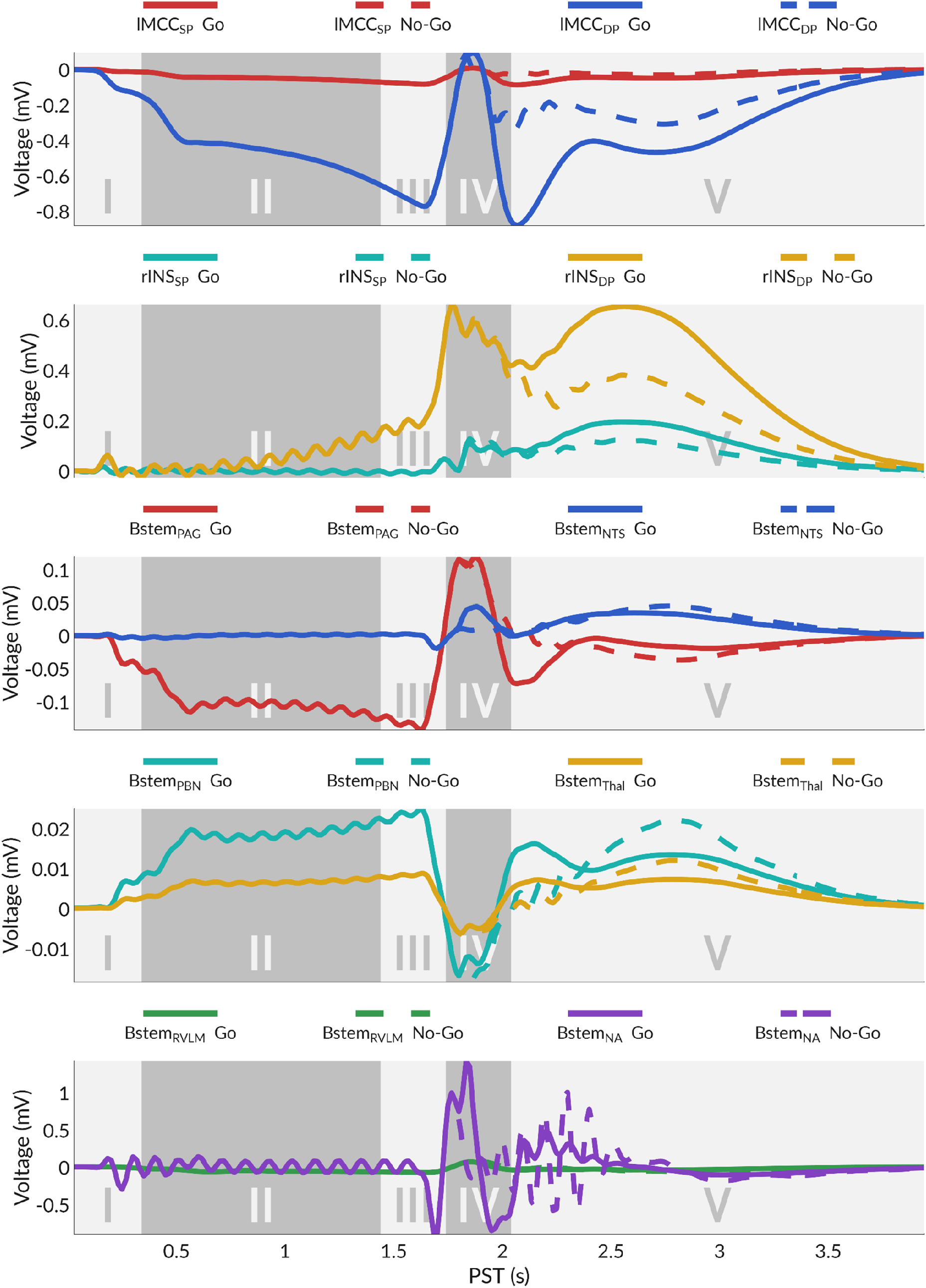
Estimated population-average membrane potentials: Posterior estimates of the population-average membrane potential for each source, comprising the viscero-sensorimotor cortical regions and brainstem nuclei. Grey shaded regions delineate the five sequential experimental phases: (I) onset cue (90-400 ms), (II) anticipation (400-1500 ms), (III) motor preparation (1500-1800 ms), (IV) target cue (1800-2100 ms), (V) motor execution (2100-4000 ms).

In the lMCC, the DP population progressively hyperpolarised from phase I through III, peaking just before target onset, while the SP population remained at baseline. In the rINS, the DP population again dominated, ramping slowly across phases I and II and depolarising sharply in phase III, with superficial layers staying quiescent throughout. Within the brainstem, the PAG showed a hyperpolarising deflection that closely followed the lMCC DP trajectory in both form and timing. The NTS showed no appreciable deviation from baseline across the same period. The PBN and thalamus both showed low-amplitude sustained depolarisations that emerged in phase I and were maintained at roughly constant levels across phases II and III. The RVLM and NA showed no sustained deviation from baseline across the anticipatory period.

### Target Cue Processing

During the target cue presentation (phase IV), a go or no-go tone was presented. The grand-averaged HR showed a transient increase following the no-go target (Figure 4C). Posterior estimates for the modulatory parameters, which capture condition-specific effects of no-go relative to go trials, are given in Table 6, with recovered trajectories in Figure 5 and membrane potentials in Figure 6; the full modulatory network is shown in Figure 7. To identify mechanisms underlying the tachycardic response, we performed BMR over the ten modulatory synaptic efficacies (Table 6), yielding 1024 reduced models (Figure 8A). As no single model won, we used BMA across the reduced space. Below we report posterior estimates from the full (unreduced) winning DCM alongside BMA.

**Figure 7.**
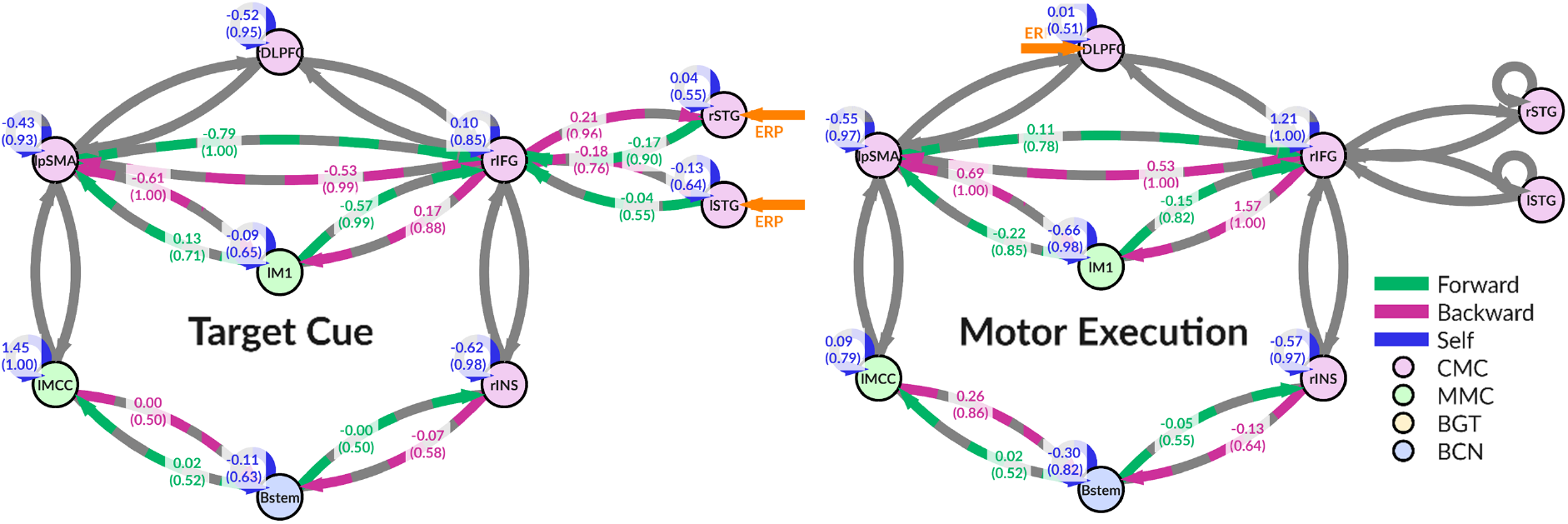
Network connectivity modulation during target cue and motor execution: No-go relative to go modulation of effective connectivity during the target cue and motor execution periods. Connection values reflect posterior expectations, with posterior probabilities in parentheses.

**Figure 8.**
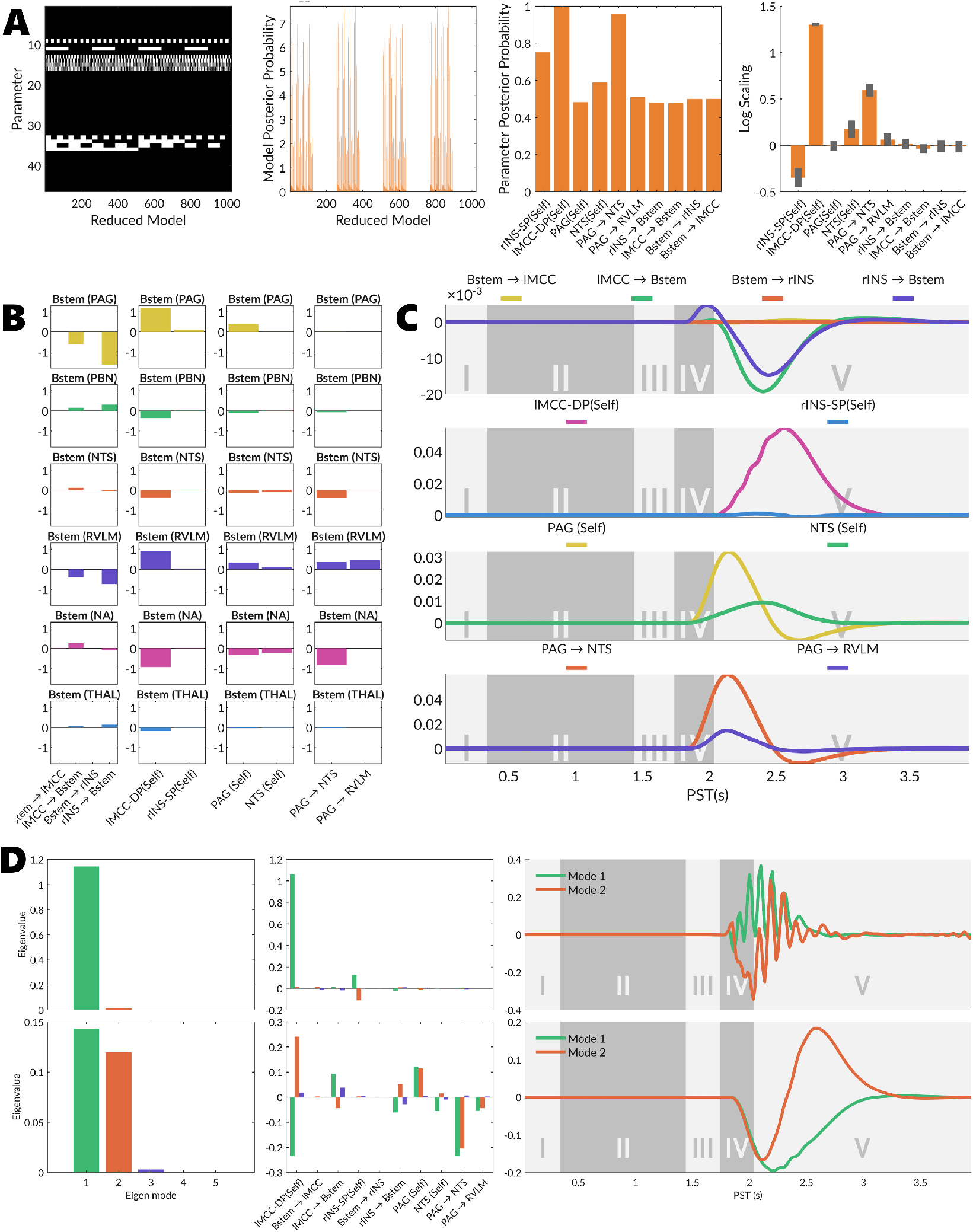
Target Cue Processing Results. **(A)** Bayesian model reduction. Left: reduced model space design (white = on, black = off). Centre-left: reduced model posterior probabilities. Centre-right: Bayesian model average parameter posterior probabilities. Right: posterior expected log scaling parameters with posterior covariance error bars. **(B)** Integrated brainstem nuclei sensitivity analysis. **(C)** HR timeseries sensitivity analysis. Grey shading delineates experimental phases I–V. **(D)** Singular value decomposition of the sensitivity of predicted EEG (top row) and HR (bottom row) responses to each parameter. Left: singular values indicating the relative contribution of each mode. Centre: parameter loadings on each mode. Right: mode time series.

**Table 6.**
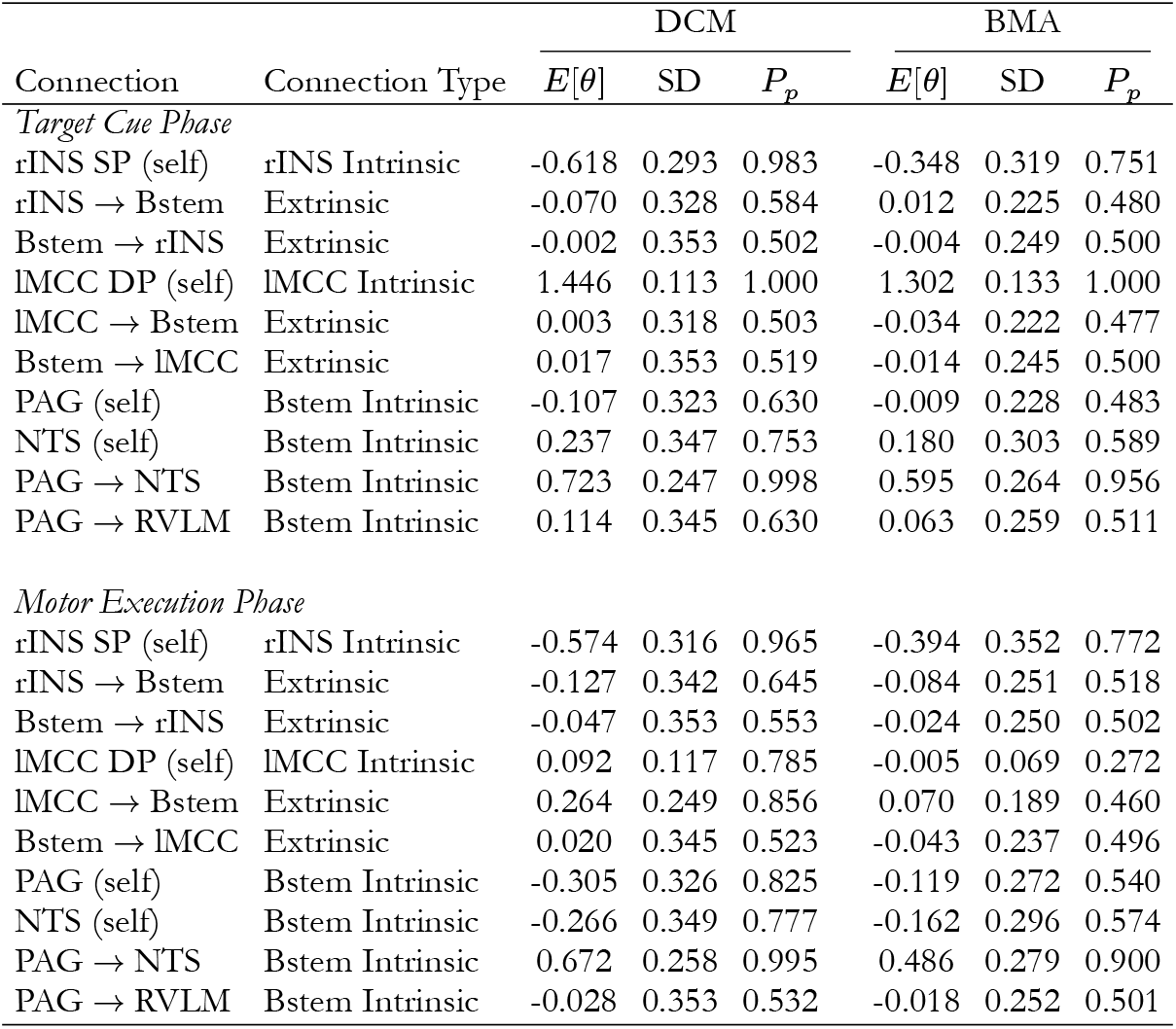
Brainstem-Related Modulatory Connections (Posterior Estimates of changes in log efficacy)

|  |  | DCM |  |  | BMA |  |  |
|---|---|---|---|---|---|---|---|
| Connection | Connection Type | *E*[*θ*] | SD | *P<sub>p</sub>* | *E*[*θ*] | SD | *P<sub>p</sub>* |
| *Target Cue Phase* |  |  |  |  |  |  |  |
| rINS SP (self) | rINS Intrinsic | -0.618 | 0.293 | 0.983 | -0.348 | 0.319 | 0.751 |
| rINS → Bstem | Extrinsic | -0.070 | 0.328 | 0.584 | 0.012 | 0.225 | 0.480 |
| Bstem → rINS | Extrinsic | -0.002 | 0.353 | 0.502 | -0.004 | 0.249 | 0.500 |
| lMCC DP (self) | lMCC Intrinsic | 1.446 | 0.113 | 1.000 | 1.302 | 0.133 | 1.000 |
| lMCC → Bstem | Extrinsic | 0.003 | 0.318 | 0.503 | -0.034 | 0.222 | 0.477 |
| Bstem → lMCC | Extrinsic | 0.017 | 0.353 | 0.519 | -0.014 | 0.245 | 0.500 |
| PAG (self) | Bstem Intrinsic | -0.107 | 0.323 | 0.630 | -0.009 | 0.228 | 0.483 |
| NTS (self) | Bstem Intrinsic | 0.237 | 0.347 | 0.753 | 0.180 | 0.303 | 0.589 |
| PAG → NTS | Bstem Intrinsic | 0.723 | 0.247 | 0.998 | 0.595 | 0.264 | 0.956 |
| PAG → RVLM | Bstem Intrinsic | 0.114 | 0.345 | 0.630 | 0.063 | 0.259 | 0.511 |
| *Motor Execution Phase* |  |  |  |  |  |  |  |
| rINS SP (self) | rINS Intrinsic | -0.574 | 0.316 | 0.965 | -0.394 | 0.352 | 0.772 |
| rINS → Bstem | Extrinsic | -0.127 | 0.342 | 0.645 | -0.084 | 0.251 | 0.518 |
| Bstem → rINS | Extrinsic | -0.047 | 0.353 | 0.553 | -0.024 | 0.250 | 0.502 |
| lMCC DP (self) | lMCC Intrinsic | 0.092 | 0.117 | 0.785 | -0.005 | 0.069 | 0.272 |
| lMCC → Bstem | Extrinsic | 0.264 | 0.249 | 0.856 | 0.070 | 0.189 | 0.460 |
| Bstem → lMCC | Extrinsic | 0.020 | 0.345 | 0.523 | -0.043 | 0.237 | 0.496 |
| PAG (self) | Bstem Intrinsic | -0.305 | 0.326 | 0.825 | -0.119 | 0.272 | 0.540 |
| NTS (self) | Bstem Intrinsic | -0.266 | 0.349 | 0.777 | -0.162 | 0.296 | 0.574 |
| PAG → NTS | Bstem Intrinsic | 0.672 | 0.258 | 0.995 | 0.486 | 0.279 | 0.900 |
| PAG → RVLM | Bstem Intrinsic | -0.028 | 0.353 | 0.532 | -0.018 | 0.252 | 0.501 |

At the cortex, the lMCC DP voltage showed a sharp depolarisation, with a condition-specific effect emerging later in the phase: Go trials produced a deep hyperpolarisation, whereas No-Go trials evoked only a shallow deflection. The lMCC DP self-inhibition showed a strong positive modulation on No-Go (DCM: 1.446, *P_p_* > 0.99; BMA: 1.302, *P_p_* > 0.99), whereas the lMCC DP to PAG and NTS connections showed no change in gain or task effect. The rINS SP voltage fluctuated modestly; its self-inhibition showed a modest reduction on No-Go that was retained in the BMA, albeit with weak posterior support (DCM: −0.618, *P_p_* = 0.983; BMA: −0.348, *P_p_* = 0.751). The rINS DP voltage carried over the prominent depolarisation from phase III, with no condition-specific effect evident. The connections between rINS and the brainstem were unremarkable, showing no clear task-dependent modulation.

In the brainstem, the PAG membrane potential sharply depolarised, closely following the trajectory of lMCC DP, with a condition-specific effect emerging late in phase IV. PAG self-inhibition decreased slightly without posterior support (DCM: −0.107, *P_p_* = 0.630; BMA: −0.009, *P_p_* = 0.483). The NTS voltage showed only modest fluctuations; its self-inhibition increased modestly with no task effect surviving BMR (DCM: 0.237, *P_p_* = 0.753; BMA: 0.180, *P_p_* = 0.589). However, the inhibitory PAG to NTS efficacy showed a prominent increase on No-Go (DCM: 0.723, *P_p_* = 0.998; BMA: 0.595, *P_p_* = 0.956). The NA, as the terminal brainstem nucleus for parasympathetic outflow, showed a sharp biphasic depolarisation at the onset of phase IV that was shorter and lower in amplitude on No-Go trials and exhibited considerably greater fluctuations than the RVLM sympathetic pathway.

We characterised the sensitivity of brainstem and cardiac responses to perturbation of synaptic efficacies, summarised as the signed area under the curve of the resulting changes in brainstem nuclei activity and as full HR timeseries (Figure 8B & C). Among the cortical-brainstem connections, descending projections dominated the responses. The rINS and lMCC projections showed similar target distributions, both reducing cumulative PAG and RVLM activity while producing small increases in PBN, NTS and thalamus, except that rINS had roughly twice the effect on PAG excitability. Both produced a biphasic HR profile comprising a brief increase during phase IV followed by a substantially larger decrease during phase V. Ascending connections exerted no appreciable influence on nuclei or HR. lMCC DP self-inhibition exerted the largest effect across the hierarchy and acted in the opposite direction to the descending projections, increasing cumulative PAG and RVLM activity while decreasing NA activity, and produced a large positive HR excursion peaking in early-to-mid phase V. By contrast, rINS SP self-inhibition was negligible for both nuclei and HR.

Among intrinsic brainstem parameters, PAG self-inhibition broadly influenced nuclei, increasing PAG, RVLM and NA activity, and produced a large HR increase with a subsequent undershoot in phase V. Of the internuclear connections, PAG to NTS efficacy had its largest effect on the NA, which it decreased, with additional increases in RVLM and a small decrease in the NTS itself, and produced an HR profile closely resembling that of PAG self-inhibition and consistent with the brief tachycardic response observed during this phase.

SVD of the sensitivity responses (Figure 8D) examined differential contributions to each modality. The electrophysiological response space was dominated by a single mode, driven almost entirely by the lMCC DP self-connection, indicating that the EEG data constrain this parameter overwhelmingly but offer little leverage over the rest. The HR response was characterised by two modes of comparable magnitude, both loading onto the same set of parameters (lMCC DP self, PAG to NTS, PAG self, lMCC to brainstem, and rINS to brainstem) distinguished by opposing signs on the lMCC DP self-connection. The second mode partially recapitulated the temporal derivative of the first mode (*R*_2_ = 0.54). The two dimensions therefore correspond approximately to the amplitude and the timing of the cardiac response, the latter arising from a temporal offset of lMCC DP self-connection perturbations relative to the other parameters (Figure 8C).

In summary, based on BMR and sensitivity analysis, the No-Go tachycardic response was principally attributed to an increase in the PAG to NTS inhibitory connection. SVD further revealed at least two distinct modes of cardiac response during this phase, corresponding to amplitude and latency.

### Motor Execution

During motor execution (phase V), participants either executed a motor response on go trials or successfully withheld a response on no-go trials. The analysis window covered an approximate two-second period encompassing both motor response and subsequent evaluative processes. Grand-averaged HR responses showed a cardiac deceleration for both conditions, but the deceleration was deeper for the no-go condition (Figure 4C). Posterior estimates for the modulatory parameters (Table 6) are reported below alongside the recovered potentials (Figure 6) and synaptic efficacy trajectories (Figure 5).

At the level of the cortex, the lMCC DP potential showed a deep sustained hyperpolarisation on Go trials that was partially suppressed during No-Go responses. Concomitantly, the condition-specific elevation in lMCC DP self-inhibition observed during phase IV for No-Go trials resolved early in phase V, such that no task-dependent differences remained by mid-phase V (DCM: 0.092, *P_p_* = 0.785; BMA: −0.005, *P_p_* = 0.272). Descending lMCC projections to the brainstem showed a positive modulation during motor inhibition that did not survive BMR (DCM: 0.264, *P_p_* = 0.856; BMA: 0.070, *P_p_* = 0.460). The rINS SP potential showed a low-level positive deflection, marginally larger on Go. The modest reduction in rINS SP self-inhibition observed in phase IV was sustained during motor inhibition (DCM: −0.574, *P_p_* = 0.965; BMA: −0.394, *P_p_* = 0.772). The rINS DP voltage showed a large sustained depolarisation peaking in early-to-mid phase V, substantially larger for Go than No-Go trials. The descending rINS projections to the PAG showed a reduction in gain, whereas those to the NTS showed little variation; neither pathway exhibited condition-specific modulation.

In the brainstem, the PAG potential recovered from its phase IV positive peak, with Go marginally more hyperpolarised before returning to baseline and No-Go showing a more gradual recovery and a shallow sustained hyperpolarisation. The NTS and RVLM potentials showed no sustained condition-specific effects. The NA showed high-amplitude oscillatory fluctuations that were larger and more prolonged on No-Go trials. The PBN recovered from the phase IV trough to a low-amplitude sustained positive deflection, with No-Go more positive in mid-phase V, and the thalamus showed a similar but lower-amplitude pattern. Among the intrinsic brainstem parameters, the PAG self-inhibition and NTS self-inhibition showed modest modulations that did not survive BMR (PAG: DCM: −0.305, *P_p_* = 0.825; BMA: −0.119, *P_p_* = 0.540; NTS: DCM: −0.266, *P_p_* = 0.777; BMA: −0.162, *P_p_* = 0.574). The PAG to NTS efficacy showed a prominent modulation that survived BMR (DCM: 0.672, *P_p_* = 0.995; BMA: 0.486, *P_p_* = 0.900), whereas the PAG to RVLM efficacy showed no clear modulation (DCM: −0.028, *P_p_* = 0.532; BMA: −0.018, *P_p_* = 0.501).

Turning to the sensitivity analysis (Figure 9B & C), the lMCC to brainstem projection had the largest effect among extrinsic connections, distributed across all nuclei. It most prominently increased cumulative NA activity, with decreases in PAG and RVLM and smaller increases in NTS and PBN, driving a large sustained negative HR excursion peaking in mid-to-late phase V. Among intrinsic connections, lMCC DP self-inhibition showed broad distributed effects, decreasing NA activity while increasing PAG and RVLM, and produced an HR increase with a late-emerging profile.

**Figure 9.**
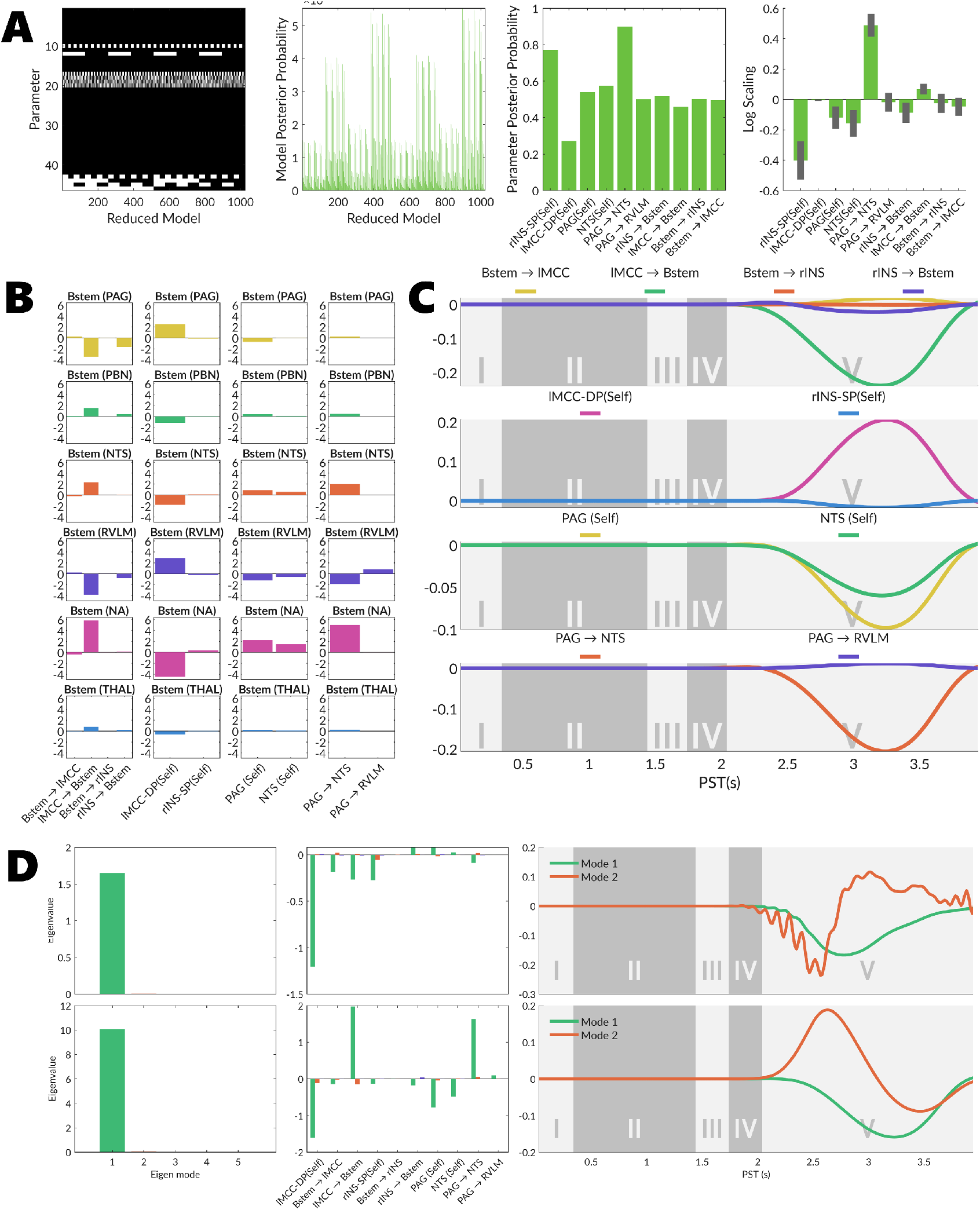
Motor Execution Results. **(A)** Bayesian model reduction. Left: reduced model space design (white = on, black = off). Centre-left: reduced model posterior probabilities. Centre-right: Bayesian model average parameter posterior probabilities. Right: posterior expected log scaling parameters with posterior covariance error bars. **(B)** Integrated brainstem nuclei sensitivity analysis. **(C)** HR timeseries sensitivity analysis. Grey shading delineates experimental phases I–V. **(D)** Singular value decomposition of the sensitivity of predicted EEG (top row) and HR (bottom row) responses to each parameter. Left: singular values indicating the relative contribution of each mode. Centre: parameter loadings on each mode. Right: mode time series.

Among intrinsic brainstem parameters, PAG self-inhibition showed a broad, modest distribution, increasing NA activity most prominently whilst decreasing PAG and RVLM, and produced a negative HR excursion with a late profile. NTS self-inhibition followed the same directional pattern as PAG and produced a smaller negative late HR excursion. Of the internuclear connections, PAG to NTS efficacy was most influential, largest at the NA, with a concomitant decrease in RVLM and increase in the NTS itself, and produced one of the largest negative HR excursions.

SVD of sensitivity responses (Figure 9D) revealed a single dominant electrophysiological mode driven over-whelmingly by lMCC DP self-connection. The cardiac response space was also characterised by a single dominant mode loaded by lMCC DP self-connection, lMCC to brainstem, and PAG to NTS efficacy, whose time course traced the slow cardiac deflection spanning mid-to-late phase V. A second mode resembling a latency component was present but of negligible magnitude, providing no evidence for a latency effect during this phase.

In summary, the PAG to NTS connection again received the strongest BMR support, and sensitivity analysis indicated that its temporal profile was concomitant with the observed deeper cardiac deceleration during motor inhibition.

## Discussion

In this work, we introduced the BCN, a neural mass model of brainstem cardiac control designed for integration within the DCM framework. This advance bridges a long-standing divide between conceptual theories of embodied and allostatic inference and the means to empirically test the functional architectures that support them. Questions around how the brain integrates peripheral physiological states and how these influence cognitive, behavioural, and affective processing can now be formally addressed in terms of functional central-peripheral neural pathways and neuronal mechanisms. This was achieved through four key advances.

The first advance provides a biophysically plausible account of brainstem nuclei and peripheral cardiac effectors, rendering the functional anatomy of cardiac regulation amenable to formal inference. The second advance extends the support to multimodal joint inversion, as demonstrated here with concurrently recorded EEG and ECG, allowing electrophysiological and cardiac data to be modelled within a single generative framework. The third advance embeds the BCN within a hierarchical, closed-loop network extending from the cortex to the periphery. This enables formal characterisation of descending commands and ascending signals across the full cortical–brainstem–cardiac axis, and how this axis interacts with other perceptual and cognitive processing streams. The fourth advance is the separation of timescales afforded by the DCM-SR framework. This means that on the faster timescale we can capture the dynamics of membrane potentials, which in turn allows us to account for how physiological states interact with ongoing perceptual processing, for instance, threat perception being biased over the cardiac cycle (Garfinkel et al. 2014). On a slower timescale, we can capture modulations in synaptic gains, thereby accounting for how physiological states can influence learning, as occurs in fear conditioning (Garfinkel et al. 2021).

In the following sections, we discuss the key findings of the present work. We first consider why the evidence was not in favour of accentuated antagonism at the SAN. We then turn to the cytoarchitectural assignments for the granular rINS and the agranular lMCC, and how their dissociable roles are concordant with our ensuing analyses. We also discuss key limitations regarding parameter identifiability, given that the BCN is silent with respect to electrophysiological observations and is principally conditioned on sluggish HR responses. We then highlight the need to address how conduction delays are handled during model inversion. We close by identifying possible extensions to the modelling repertoire to further expand the sorts of observations and questions that can be addressed.

### Accentuated Antagonism

The decisive support for the linear SAN model indicates that the present data did not require the inclusion of nonlinear accentuated antagonism, the potentiation of vagal slowing that occurs when sympathetic tone is concurrently elevated. On the face of it, this may be unexpected given that no-go target presentation elicited a brief tachycardic response, followed by a deepening of cardiac deceleration during motor inhibition. The observation of this biphasic pattern might be taken to constitute the necessary conditions for accentuated antagonism. Specifically, it could suggest sympathetic activation closely succeeded by parasympathetic recruitment. However, the analyses instead pointed to a parasympathetically mediated response, arising from modulation of the vagal brake via the PAG to NTS pathway operating on a linear model of SAN innervation, as providing a more parsimonious account of the observed dynamics. Even granting a sympathetic contribution to the observed responses, it would likely be too small and too brief to shift the operating point onto the steep, gain-amplifying region of the bivariate autonomic surface. Consistent with this interpretation, the supporting literature on accentuated antagonism has relied predominantly on paradigms with robust autonomic perturbations such as postural (Uijtdehaage and Thayer 2000) or pharmacologically induced stress (Tulppo et al. 1998), with little precedent for cognitive tasks of the kind employed here. Nevertheless, the capacity to characterise such nonlinear regimes remains accessible within the BCN model, and it will be interesting to determine whether evidence of accentuated antagonism can be found under clearer-cut sympathetic-provoking conditions, such as stress-or threat-inducing experimental designs.

### Allostatic Inference Functional Cytoarchitecture

Under predictive processing and active inference, the laminar cytoarchitecture of a cortical region is proposed to map onto its functional orientation within hierarchical message passing, with granular regions subserving ascending sensory signalling and agranular regions issuing descending motor predictions (Shipp, Adams, and Friston 2013; Bastos et al. 2012). Barrett and colleagues extended this principle to interoception, proposing that granular cortices receive ascending interoceptive prediction errors whilst agranular visceromotor cortices issue allostatic commands to brainstem effectors (Barrett and Simmons 2015; Barrett, Quigley, and Hamilton 2016). While this account has received empirical support via DCM analysis in somato-sensorimotor domains (Bhatt et al. 2016; van Wijk et al. 2018), its application to viscero-sensorimotor processing has not been formally tested. The present work therefore provides an initial evaluation of competing cytoarchitectural assignments for viscero-sensorimotor cortical regions within a generative modelling framework.

In the present work, BMC identified the lMCC as best represented by an agranular microcircuit and the rINS by a granular microcircuit, suggesting that the lMCC issues descending allostatic predictions whilst the rINS receives ascending interoceptive prediction errors. These assignments should be read as functional rather than structural: they do not constitute claims about the laminar composition of either region, but indicate that, under the present paradigm, the rINS operated principally as a recipient of ascending interoceptive signals rather than as a source of descending visceromotor commands. These assignments are consistent with prior evidence localising effort-related cardiovascular arousal to the MCC, both in lesion studies, where damage to this region blunts autonomic responses during effortful task performance (Critchley et al. 2003), and in intracranial recordings, where MCC activity drives anticipatory HR modulation (Gillies et al. 2019). The agranular assignment of the lMCC thus positions it well to generate descending allostatic predictions concerned with upcoming action demands. The granular assignment of the rINS, by contrast, aligns with its established role in representing internal bodily states (A. D. Craig 2009) and tracking the accuracy of cardiac interoceptive judgements (Critchley et al. 2004). Additionally, its engagement during conscious error perception and salience monitoring points to a broader evaluative function, one that registers (precision-weighted) prediction errors arising from both interoceptive and exteroceptive sources, consistent with its position as a hub for integrating viscero- and somato-sensory salience within the context of task performance (Ullsperger et al. 2010).

Beyond the cytoarchitectural assignments, the generative DCM framework affords deeper insight into cortical-brainstem-cardiac hierarchical message passing. In the section that follows, we illustrate the explanatory capacity of this framework by showing its ability to account for message passing between cortical and brainstem regions and for intrinsic brainstem internuclear processes, and we furnish these findings with an active inference interpretation of cardiac allostasis and interoception.

### Target Cue Processing

We first revisit the architecture of the BCN to understand candidate mechanisms that can produce the tachycardic event seen following No-Go target presentation. The model assumes a default parasympathetic policy favouring bradycardia, which is principally changed via PAG excitation to engage active coping strategies. The BCN simplifies the PAG’s competing multi-columnar architecture by treating its functional state as a single axis, where depolarisation promotes active coping policies, and hyperpolarisation reinforces default quiescent policies. Active policies can then manifest as either inhibition of the NTS (withdrawal of the vagal brake) or direct excitation of the RVLM (direct sympathetic drive).

During the anticipatory period preceding target presentation, the lMCC DP potential became progressively hyperpolarised relative to baseline. Viewed as a descending influence on the PAG, this sustained hyperpolarisation deepens the default parasympathetic policy, and offers a plausible mechanism for the anticipatory cardiac deceleration observed over this window.

Following target presentation, the lMCC DP exhibited a brief depolarising potential that can be interpreted as a shift towards active coping policies. Concomitantly, lMCC DP self-inhibition was substantially increased during No-Go presentation, resulting in a shallower afterhyperpolarisation potential that emerged late in target cue processing and became more pronounced during motor response. Sensitivity analyses indicated the capacity of lMCC DP self-inhibition to drive a tachycardic response, although its temporal profile emerged considerably later than the observed HR changes. Thus, whilst increased lMCC DP self-inhibition can reasonably be interpreted as attenuation of a descending motor command for an expected action policy, its contribution to the tachycardic event remains uncertain.

In contrast, our findings support the tachycardic event being driven by the inhibitory PAG to NTS pathway. This connection showed a substantial increase during No-Go presentation, and given that the NTS constitutes the primary source of parasympathetic control, augmentation of this inhibitory influence is consistent with PAG-mediated vagal brake withdrawal. Sensitivity analyses yielded a temporal profile closely matching the observed HR changes, supporting its role as the most likely tachycardic mechanism.

We also found weaker evidentiary support for a reduction in rINS SP self-inhibition during No-Go presentation, an effect that may index an upweighting of ascending interoceptive prediction errors. If No-Go entails attenuation of the expected action policy, the resulting tachycardic episode constitutes a deviation from the predicted bodily state, and increased precision of ascending interoceptive signals would be expected to refine subsequent predictions. As such, the rINS effect, whilst tentative, aligns with a broader picture in which the hierarchy shifts towards heightened interoceptive updating during withholding of a prepared action.

### Motor Response

During the motor response period, both conditions elicited a transient cardiac deceleration that was deeper on No-Go trials. This phase revealed condition-specific membrane potential effects that had been only weakly evident during target cue processing. At the cortical level, lMCC DP was less hyperpolarised on No-Go trials, whilst rINS DP was less depolarised on No-Go compared to Go. At the brainstem level, the PAG also differed between conditions; Go trials showed a brief hyperpolarisation followed by a rapid return to baseline, whereas No-Go trials showed a gradual reduction that entered into a shallow but prolonged hyperpolarisation.

The increased gain of the inhibitory PAG to NTS connection carried over into this period. Interestingly, sensitivity analysis now showed that this connection could account for the deeper bradycardia observed during No-Go trials, whereas previously it was responsible for driving the tachycardia. Tentatively, this can be explained by the shallow hyperpolarised state of the PAG during No-Go trials, which may be interpreted as a switch from an active to a passive coping strategy, thereby disinhibiting the NTS. The lMCC to brainstem projection similarly produced a profile that could plausibly account for the deeper bradycardia on No-Go trials, though it lacked strong BMR support. However, its heavy loading in the SVD of cardiac responses points to a potential parameter identifiability issue, which we address below.

BMR again sustained weak support for reduced rINS SP self-inhibition during No-Go motor execution, which may suggest that interoceptive errors were afforded increased precision during motor response inhibition and evaluation.

### Parameter Identifiability

An important consideration when deploying the BCN is the identifiability of brainstem states and parameters. The BCN was treated as silent towards electrophysiological observations, meaning parameters were likely mostly constrained by cardiac observations as they were most proximal. While precedent exists for such silent nodes in DCM studies (David et al. 2011), this case is distinct because HR responses are comparatively sluggish and therefore act as a low-pass filter on brainstem latent state dynamics. This places the BCN in a similar setting to fMRI modelling (K. J. Friston et al. 2000), which has comparatively fewer temporal modes, suggesting that the number of parameters estimated from cardiac responses should be treated with caution.

In the present work, we found a maximum of two temporal modes, which corresponded to the amplitude and latency of the cardiac response. This is consistent with the sensitivity analysis, which showed that different parameters can produce near-identical temporal profiles. This indicates that one needs to be selective about which parameters are left free to vary. It should be noted that the paradigm employed here was relatively flat in its interoceptive and allostatic demands. As such, this question of identifiability should be considered an open question and addressed with paradigms employing stronger autonomic engagement and interoceptive evaluative demands. Model comparison and BMR were also performed on a single grand-averaged DCM under fixed effects; the evidence therefore reflects the group-mean response and does not account for between-participant variability, which a hierarchical extension using parametric empirical Bayes across participants would address.

### Limitations

A key limitation of the present work concerns the handling of conduction delays within the inversion scheme. The current implementation integrates using spm_int_L, which incorporates conduction delays via a firstorder Taylor approximation rather than explicitly tracking delayed state histories. This incurs non-negligible integration error when delays exceed roughly 10% of the intrinsic time constants of the modelled system (Lemaréchal, George, and David 2018; Schöbi et al. 2021). For the cardiac pathways in the BCN, this is a serious concern. Vagal efferent latencies at the SAN are of the order of 50-100 ms (Jalife et al. 1983), sympathetic efferent delays extend to 400-700 ms due to unmyelinated C-fibre conduction and noradrenergic kinetics (Mokrane and Nadeau 1998), and the ascending baroreceptor arc contributes a further 500-600 ms (Borst and Karemaker 1983).

Alternative delay-compatible integrators exist, notably the LDE scheme (Schöbi et al. 2021) and the dde23-based approach (Lemaréchal et al. 2022), but both carry substantially increased computational cost, even for standard ERP epochs. The BCN operates over considerably longer trial windows, and it remains unclear whether these schemes remain tractable at that scale. A more practical route may be to use the high-order Taylor approximation described in Appendix A of Karl J. Friston et al. (2014) to accommodate long peripheral delays without incurring the full overhead of explicit delay-differential equation solvers.

Without accurately accounting for these delays, the temporal ordering of ascending and descending signals cannot be correctly identified. This is particularly problematic for arbitrating between baroreceptor and proprioceptive accounts of cardiac timing inference, as each pathway carries a distinct delay profile (Benarroch 1993; Al et al. 2023; A. D. Levy, Zeidman, and Friston 2026b). Resolving this is also necessary for testing the bidirectional interactions between cardiac and perception-action cycles (Al et al. 2023; Palser et al. 2021; Galvez-Pol, McConnell, and Kilner 2020; Skora, Livermore, and Roelofs 2022; Allen et al. 2022). Until this is resolved, parameter estimates for delays and the temporal ordering of ascending and descending message passing should be interpreted with caution.

### Future Work

Near-term work will focus on further validating the BCN model. Experimental paradigms with greater allostatic and interoceptive demands will be essential to further illustrate the capacity to arbitrate competing anatomical hypotheses. Pharmacological manipulations using agents with peripheral and central specificity, such as propranolol, atropine, and non-brain-penetrant compounds like methylatropine, will be particularly valuable for establishing that the BCN can dissociate central from peripheral mechanisms. Complementing these approaches, optically pumped magnetometers offer a promising route to provide more direct and constrained observations of brainstem electrophysiological responses (Tierney et al. 2021).

Looking further ahead, the framework opens the door to addressing broader questions of embodied inference. The framework was designed to readily accommodate other peripheral systems. With the inclusion of a wider repertoire of generative models of visceral and volitional systems, formal testing of embodied inference becomes tractable. This capacity to embed multiple peripheral models within a single hierarchical framework enables formal tests of how the brain coordinates competing allostatic and perceptual demands. For example, by incorporating a generative model of the oculomotor system alongside cardiac control, one could test how the planning of saccadic eye movements recruits cardioballistic facilitation, alongside concomitant cardiac deceleration to aid visual perceptual processing.

### Conclusion

We have introduced the BCN, the first generative model of brainstem cardiac control designed for formal inversion against multimodal neural and cardiac data within the DCM framework. We demonstrated its ability to adjudicate between granular and agranular cytoarchitectural assignments for viscero-sensorimotor cortical regions, and to characterise the respective contributions of descending visceromotor commands and ascending interoceptive signals to cardiac regulation during motor inhibition. By rendering the functional architecture of brain–heart communication amenable to formal inference, the BCN offers a framework for characterising how disruptions in the balance between descending allostatic predictions and ascending interoceptive evidence contribute to disorders of interoception and autonomic regulation, such as anxiety and somatic symptom disorders. More broadly, this provides a route to mechanism-based stratification and targeted intervention by dissociating central from peripheral contributions to autonomic dysregulation.

